# Environmental dynamics shape human learning: change points versus random walks

**DOI:** 10.1101/2025.11.26.690700

**Authors:** Cedric Foucault, Lilian A. Weber, Laurence Hunt

## Abstract

Adaptive learning requires interpreting prediction errors in light of environmental dynamics. However, environments may differ not only in *when* changes occur, but also in *how* they occur: some nonstationary processes drift gradually, whereas others change abruptly. This raises the question of whether humans can adapt their learning to the generative structure of their environment. Here, we compared learning in two canonical nonstationary environments: abrupt change points versus gradual random walks. Across two behavioral experiments using a predictive inference task and a unified Bayesian framework, we show that participants adjust their learning to the dynamics of their environment. Cross-validated model comparison showed that the environment-matched model best explained behavior in nearly all participants. Human apparent learning rates also showed the predicted dynamics-specific signature, increasing more sharply with prediction-error magnitude under change points compared to random walks. Participants tracked the changing latent variance (environmental stochasticity) online in both environments, jointly with the latent mean, through coordinated adjustments of paddle width and location. In the change-point environment, their width adjustments further distinguished changes in variance from changes in mean. Together, these results show that humans adapt their learning to both how latent states change and which properties are changing. They establish a unified computational account of adaptive learning across different environmental dynamics.

## Introduction

Human learning and decision-making often take place in dynamic, uncertain environments. A key computational challenge is to infer latent properties of the environment—such as means and variances—from noisy observations, and to adapt to potential changes in these properties over time.

For example, consider a commuter noticing that their travel time is longer than expected. To decide whether to leave earlier or take a different route on subsequent days, they must first judge whether this fluctuation is simply part of the typical day-to-day variability (the variance), or whether there has been a genuine change in travel conditions. If a change is likely, they must further consider the nature of the change: is it a gradual increase in traffic (e.g., due to a growing local population), or a sudden disruption (e.g., new roadworks)? Effective decision-making therefore requires inferring both the mean and the variance of travel times, and adapting based on whether these properties are stable, gradually shifting, or abruptly changing.

Two canonical classes of dynamics are commonly used to describe how latent variables evolve over time: *change-point dynamics*, in which the latent variable is assumed to change abruptly and discretely (Adams & MacKay, 2007; Behrens et al., 2007; Nassar et al., 2010), and *random-walk dynamics*, in which the latent variable drifts gradually and continuously (Kalman, 1960; Mathys et al., 2011; Piray & Daw, 2021; Pulcu & Browning, 2025). Both frameworks capture essential aspects of environmental dynamics but fundamentally differ in how they represent environmental changes. These differences, in turn, imply distinct statistical and computational principles for learning.

Are humans able to learn appropriately given the environment they are placed in? A large body of work has examined human learning in either change-point environments (e.g., (Behrens et al., 2007; Glaze et al., 2015; Vaghi et al., 2017)) or random-walk environments (e.g., (Daw et al., 2006; Lee et al., 2020; Piray & Daw, 2024; Pulcu & Browning, 2025)) (for an overview of major studies and their environmental dynamics, see also Table 1 in (Foucault, 2023)). These two types of environments have largely been investigated separately. One exception is (Nassar et al., 2019), who compared a change-point condition with an oddball condition whose latent mean followed a random walk; (Razmi & Nassar, 2022) subsequently modeled learning across these contexts. However, neither study formally compared how well Bayesian models assuming change-point versus random-walk mean dynamics predicted individual participants’ responses in each environment.

It remains an open question whether participants apply the appropriate model to the appropriate environment. This gap in our understanding of human learning is perhaps due to a lack of consistent study designs, task structures, analysis methods, and computational models across the two largely parallel lines of research—one anchored in change-point models and the other in random-walk models. Studies in change-point environments typically adopt a change-point modeling framework, analyzing modulations in learning rate in terms of variables such as the estimated change-point probability (Foucault & Meyniel, 2024; McGuire et al., 2014; Murphy et al., 2021; Nassar et al., 2012; Vaghi et al., 2017). Conversely, studies in random-walk environments typically adopt a random-walk modeling framework, relating human learning rates to model-derived estimates of volatility—that is, the diffusion noise of the random walk—(Mathys et al., 2011; Piray & Daw, 2021; Pulcu & Browning, 2025). Although some studies have analyzed human data from both types of environments, they have generally relied on only a single model class (typically a random-walk model) to interpret behavior (Mathys et al., 2011; Piray & Daw, 2020, 2021). As a result, it is unclear whether findings attributed to one model class might alternatively be explained by the other. This has important implications beyond basic learning research: work in computational psychiatry has shown that reanalyzing existing datasets under alternative generative-model assumptions can yield different interpretations of the computational mechanisms underlying psychiatric traits and symptoms (Mikus et al., 2025; Piray & Daw, 2021). Theoretical work on simulated data has shown that change-point and random-walk models could be disambiguated (Marković & Kiebel, 2016), but whether humans align more closely with one model or the other remains to be determined.

A further outstanding question concerns whether and how humans infer and respond to changes in variance (stochasticity) over time within a sequence, alongside changes in the mean. Most previous studies either did not manipulate variance, or if they did, varied it across experimental blocks—keeping it fixed within blocks (Lee et al., 2020; McGuire et al., 2014; Nassar et al., 2010, 2012; Piray & Daw, 2024). Yet in many natural environments, both the central tendency and the variability—which determines the reliability of individual observations—can change, creating the added challenge of distinguishing changes in mean from changes in variance. This is non-trivial because both types of changes can produce a similar increase in the dispersion of recent observations and in prediction errors. Thus, an open question is whether humans can infer a changing variance alongside a changing mean, distinguish between change points in mean versus variance, and explicitly report when they believe the variance has changed. Being able to report such changes is important, for example because communicating uncertainty can substantially improve decision-making (Bahrami et al., 2010; Van Der Bles et al., 2019). Moreover, the way in which learners infer variance may interact with how they infer the mean. These interactions may differ depending on whether the underlying dynamics follow a change-point or random-walk process.

Overall, the influence of environmental dynamics on human learning remains poorly understood. In particular, it is unclear whether humans learn differently and whether they deploy the appropriate model in environments governed by change points versus random walks, and how these differences may be captured by computational models. A direct comparison of the two types of environments and their corresponding models—under conditions where both mean and variance can change—is currently lacking.

Here, we directly compare human learning under change-point and random-walk dynamics with the predictions of Bayesian models, within a single task that requires joint inference of mean and variance. Across two experiments, participants performed a learning task in which the generative mean changed through either abrupt change points (Experiment 1) or a gradual random walk (Experiment 2), while the generative variance switched between two states. We analyze behavior using model-agnostic measures derived directly from participants’ responses and a unified family of Bayesian models with alternative generative-model assumptions. We formally compare these models through participant-level model fitting and cross-validation, evaluating their predictive performance on held-out behavioral data. We complement these comparisons with simulations from the fitted models, assessing how well the models reproduce qualitative behavioral signatures. Our results show that the environment-matched model best explained behavior in nearly all participants. Human apparent learning rates also showed the predicted dynamics-specific signature, increasing more sharply with prediction-error magnitude under change-point than random-walk dynamics. Moreover, participants’ paddle-width choices tracked changes in generative variance online in both environments; in the change-point environment, these choices further distinguished changes in variance from changes in mean. A graded hybrid extension of the model family, fitted to human data, also showed a strong environment-dependent shift in the relative weighting of the change-point and random-walk predictions. Together, these findings show that humans adapt their learning to both how latent states change and which properties are changing, providing a unified computational account of adaptive learning across environmental dynamics.

## Results

### Task structure and performance

We designed a learning task and conducted two experiments to examine how environmental dynamics influence human learning (Figure 1). In both experiments, the environment contained a generative mean and a generative variance that independently changed over time, and participants’ ability to infer them online simultaneously was directly tested by the task.

**Figure 1.**
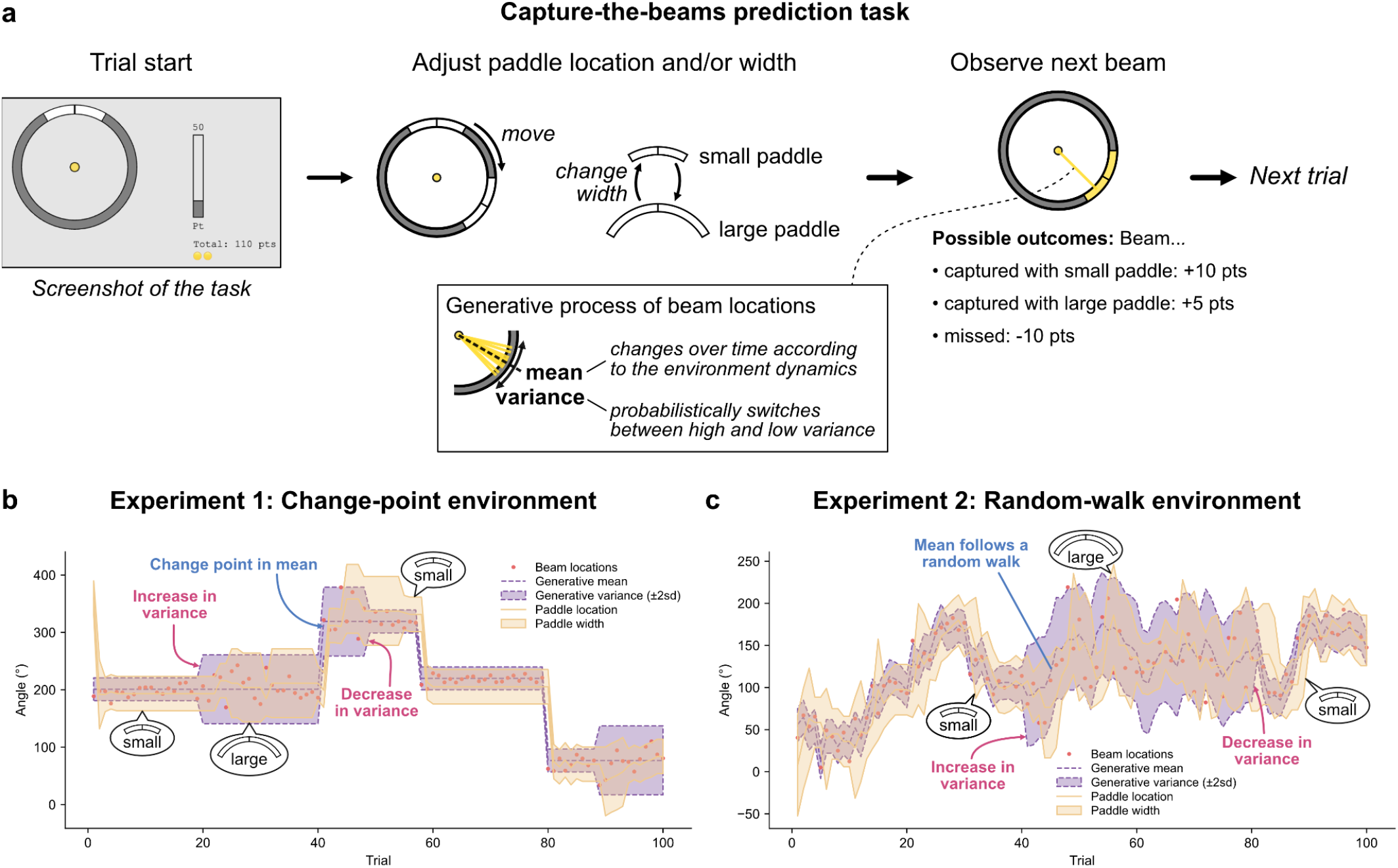
Capture-the-beams prediction task. (a) Task structure. On each trial, participants positioned a paddle on a circular ring and selected its width (small or large) to predict the angular location of the next beam. After adjusting the paddle, they observed the next beam and received feedback. Capturing a beam with the small paddle yielded more points (+10) than with the large paddle (+5), but carried a higher risk of misses, which incurred a loss (–10). Beam locations were generated from a latent mean and variance that both changed over time. In Experiment 1, the generative mean changed abruptly at unpredictable change points, whereas in Experiment 2, it changed gradually and continuously from trial to trial according to a random walk. In both experiments, the generative variance alternated probabilistically between low- and high-variance states. Participants: n = 30 in each of the two experiments. (b, c) Example sequences from Experiment 1 (change-point environment) and Experiment 2 (random-walk environment), and example participant, whose paddle locations and widths (orange) tracked the generative mean and variance (purple), respectively.

On each trial, participants predicted the next observation in a sequence—corresponding in this task to the beam emission angle around a central point (Figure 1a). Instead of a single-point prediction, participants provided a prediction interval by adjusting a *paddle*, whose angular extent defined the interval. They could rotate the paddle around a circular ring and choose between two available paddle sizes (small or large). Using the small paddle resulted in higher gains if the next beam was captured but carried a higher risk of missing the beam, which induced a loss. Once they confirmed their choice, the next beam appeared, the resulting gains or losses were displayed, and the next trial began.

Accurate performance required learning, through inference, two latent variables: the *generative mean* and the *generative variance*, which governed the distribution from which the next beam emission angle was drawn. Importantly, the optimal paddle location and size directly reflected these inferences. To maximize their score, participants needed to center the paddle on the current generative mean, and choose the large paddle when the generative variance was high or the small paddle when the generative variance was low (see Methods). This mapping provided a trial-by-trial behavioral readout of participants’ beliefs about both latent variables.

Two experiments were conducted with 30 participants each. In Experiment 1, the dynamics of the generative mean followed a change-point process (Figure 1b), whereas in Experiment 2, they followed a random-walk process (Figure 1c). Apart from the dynamics of the generative mean, all other task parameters were identical between the two experiments.

To make the task intuitive, we embedded it in a space-explorer cover story in which participants captured energy from beams emitted by a distant star to power their colony. We explained the key components of the generative process using analogies. We first introduced the generative mean and variance using a flashlight analogy: the star’s average emission direction represented the mean, and the spread of the beams the variance. We then introduced the two variance levels as two distinct states of the star—a focused state (low variance) and a diffuse state (high variance). Lastly, we explicitly described how the generative variables changed: participants in Experiment 1 were told that the star’s average direction changed abruptly from time to time, whereas participants in Experiment 2 were told that it drifted gradually; both groups were told that the star occasionally switched between focused and diffuse states. Parameter values, including the frequency and magnitude of changes and the precise variance levels, were not disclosed. Before the task began, we administered a short comprehension quiz to ensure that participants had understood these concepts and their implications for maximizing score.

Participants performed the tasks accurately: Experiment 1, percentage of beams captured 81 ± 1%, proportion of trials using the small/large paddle 66/34% ± 3%, score 586 ± 9 points per 100-trial block; Experiment 2, beams captured 77 ± 1%, small/large paddle 54/46% ± 5%, score 447 ± 11 points (values are mean ± s.e.m.).

### Computational modeling framework

To derive theoretical predictions about behavior in our task, we developed and simulated a set of computational models (Figure 2). These models formalize specific hypotheses about the inference processes humans might use, allowing us to predict how behavior should unfold under different assumptions and, by comparison with human data, gain insights into the computational processes underlying human learning and decision-making.

**Figure 2.**
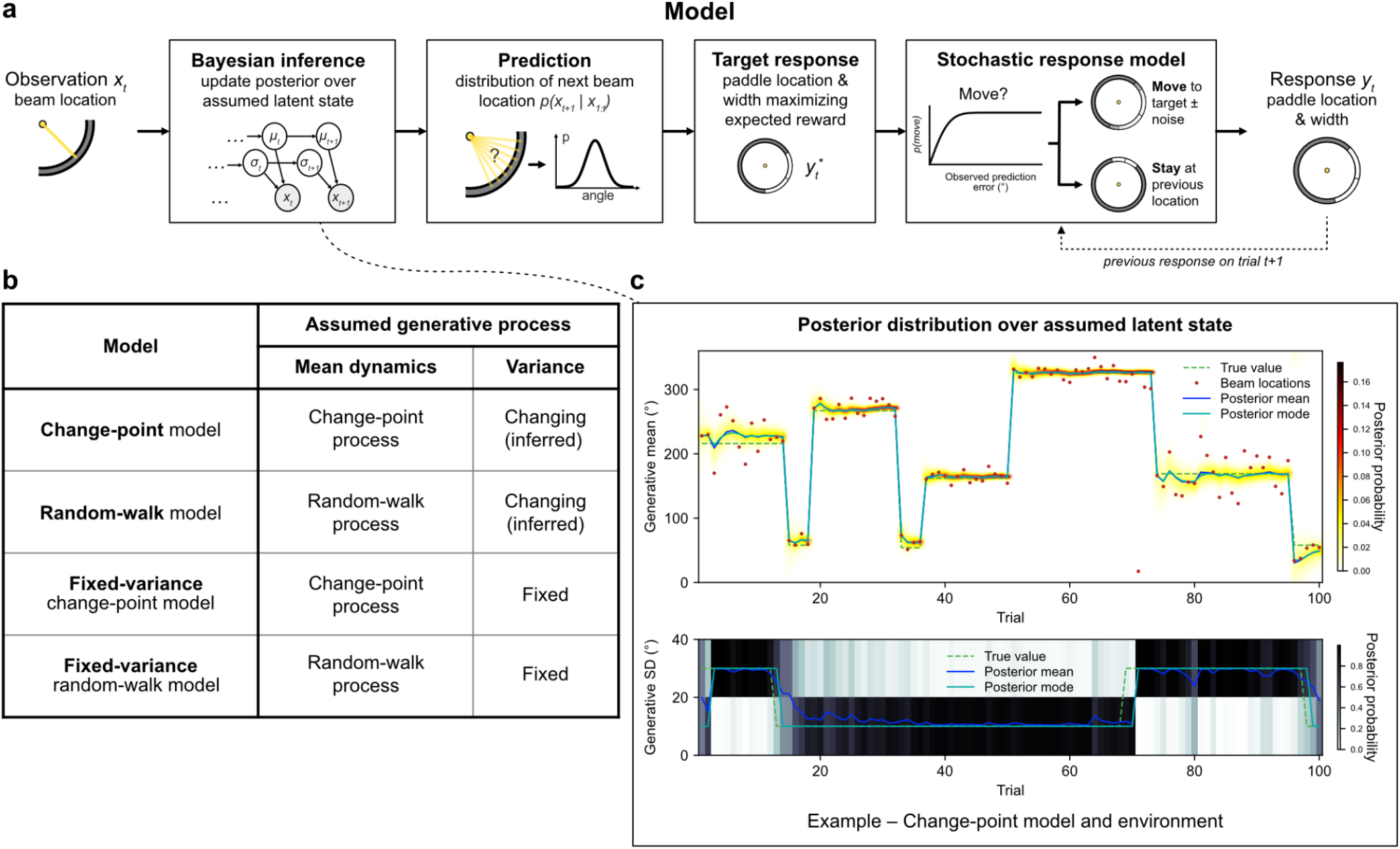
Modeling framework. (a) Model architecture. On each trial, a new beam location *x_t_* is observed. Through Bayesian inference, the model updates its posterior over its assumed latent state. From this posterior, the model derives the predictive distribution for the next beam location and determines the target response, *y_t_\**—the paddle location and width that maximize expected reward. The stochastic response model then generates the actual response, *y_t_*, through a probabilistic movement decision: movement probability increases with the observed prediction error (defined as the circular distance between the observed beam and the previous paddle location). If movement is initiated, the paddle is moved to the target location with angular response noise; otherwise, it stays at its previous location. In both branches, an error probability applies to the paddle-width choice. (b) Model family. Models differ in their assumed dynamics of the generative mean (change-point or random-walk process) and whether generative variance changes and is inferred or is fixed. See the main text for how models proposed in prior work relate to this model space. (c) Example posterior inference for the change-point model in the change-point environment. Heat maps show the marginal posterior distributions over the generative mean (top) and the generative variance, represented by its standard deviation (bottom). Posterior means and modes are overlaid with the true latent values; observed beam locations are shown in the top panel.

Our model set was defined in a systematic way to isolate the role of key features of the assumed generative process (Figure 2b). In particular, we varied (i) the type of dynamics (abrupt change points versus gradual random walks) and (ii) whether the generative variance was assumed to be fixed or changing. This yielded a family of models that can be compared and contrasted to provide theoretical insights.

Several influential models from prior work can be located within this model space. These correspondences concern assumptions about the generative process rather than identical inference algorithms. Our formulation unifies these related generative assumptions within a common Bayesian framework. The fixed-variance change-point model makes the same core generative assumptions as the models used in (McGuire et al., 2014; Nassar et al., 2010, 2012; Vaghi et al., 2017). The fixed-variance random-walk model corresponds to classical Kalman-filter formulations and their applications (Daw et al., 2006; Kalman, 1960; Lee et al., 2020). The closest precedents for the random-walk model, which infers changing variance, are models that jointly estimate stochasticity and volatility (Piray & Daw, 2021, 2024; Pulcu & Browning, 2025), although these differ in how their variance parameters evolve and how inference is implemented. For the remaining cell—the change-point model, which also infers changing variance—none of the works cited here have the same generative structure. A reduced-model variant in (Nassar et al., 2010) inferred an unknown observation variance, but this variance was stationary, whereas our models with variance inference assume that it changes over time.

After observing a new beam location, *x_t_*, each model generates a response through four stages (Figure 2a).

**(1) Bayesian inference:** The model updates its posterior over the latent state under its assumed generative process. When the assumed generative process matches the environment, the posterior represents Bayes-optimal inference and closely tracks the true latent state, as illustrated in Figure 2c.
**(2) Prediction:** From the updated posterior, the model derives the predictive distribution over the next beam location.
**(3) Target response:** The model determines the paddle location and width that maximize expected reward under this predictive distribution. These define the model’s target response, *y_t_\**.
**(4) Stochastic response model:** The stochastic response model generates the actual response, *y_t_*, from the target response, *y_t_\**. It comprises a movement-initiation decision—whether to stay at the previous paddle location or move—followed by generation of the paddle location and width conditioned on this decision. The movement-initiation probability increases with the observed prediction error—the circular distance between the current beam location and the previous paddle location. If movement is initiated, the model generates a paddle location centered on the target location with angular response noise; otherwise, the paddle remains at its previous location. In either branch, the paddle-width choice is subject to an error probability.

The stochastic response model allows small prediction errors to result in no paddle movement, a tendency documented in belief-updating tasks that may reflect response costs or a judgment that the current response is “good enough” (Bruckner, Nassar, et al., 2025; Gershman, 2020; Nassar et al., 2021; Simon, 1956). It also provides the probabilistic link needed to fit each inference model to participants’ trial-by-trial responses and accommodate individual differences in movement initiation, angular response variability, and width-choice errors through parameters estimated separately for each participant. The response model is therefore a descriptive and non-normative component of the full model, which is separate from the inference component to which all normative claims are restricted.

### Model predictions: adaptation to environmental dynamics and stochasticity

Having developed this family of models, we next examine their predictions about behavior under different environmental dynamics (change points vs. random walks). To characterize learning behavior in a model-agnostic manner, we quantified an *apparent learning rate*—defined on each trial as the signed update in paddle location divided by the signed prediction error (Foucault & Meyniel, 2024; Nassar et al., 2010). This measure is not a fitted model parameter but a descriptive index of how strongly an agent updates its response to new evidence. For simplicity, we refer to it below as the *learning rate*.

Model simulations show that the environmental dynamics dictate how this learning rate should vary with the observed prediction error: the model adapts its learning rate markedly differently when assuming change-point dynamics compared to random-walk dynamics (Figure 3). In the change-point model, the learning rate increases sharply with prediction error, reaching values close to 1 for large errors (Figure 3a). By contrast, in the random-walk model, the learning rate shows little or no modulation once beyond the low-error regime: increases in the magnitude of the prediction error do not translate into larger learning rates (Figure 3b). This qualitative difference reflects the underlying assumptions about environmental dynamics. In a change-point process, a large prediction error provides evidence that a change point in the latent mean may have occurred, in which case it is optimal to reset the belief about the latent mean and weight the new observation heavily—hence the steep rise in learning rate for large errors. Conversely, when prediction errors are small to moderate (e.g. < 50°), a change point is unlikely, implying that the latent mean has remained stable; in this case, the model exhibits a very low learning rate. In a random-walk process, however, the latent mean is assumed to drift gradually on every trial rather than change abruptly, thus, belief updates should be applied consistently (even when prediction errors are only moderate in size). As a result, the random-walk model exhibits a relatively stable learning rate across different error magnitudes, while the change-point model flexibly suppresses or amplifies updates depending on the inferred probability of a change point. This predicted difference between the models was preserved across a broad range of response-model parameter values (Supplementary Figure 1).

**Figure 3.**
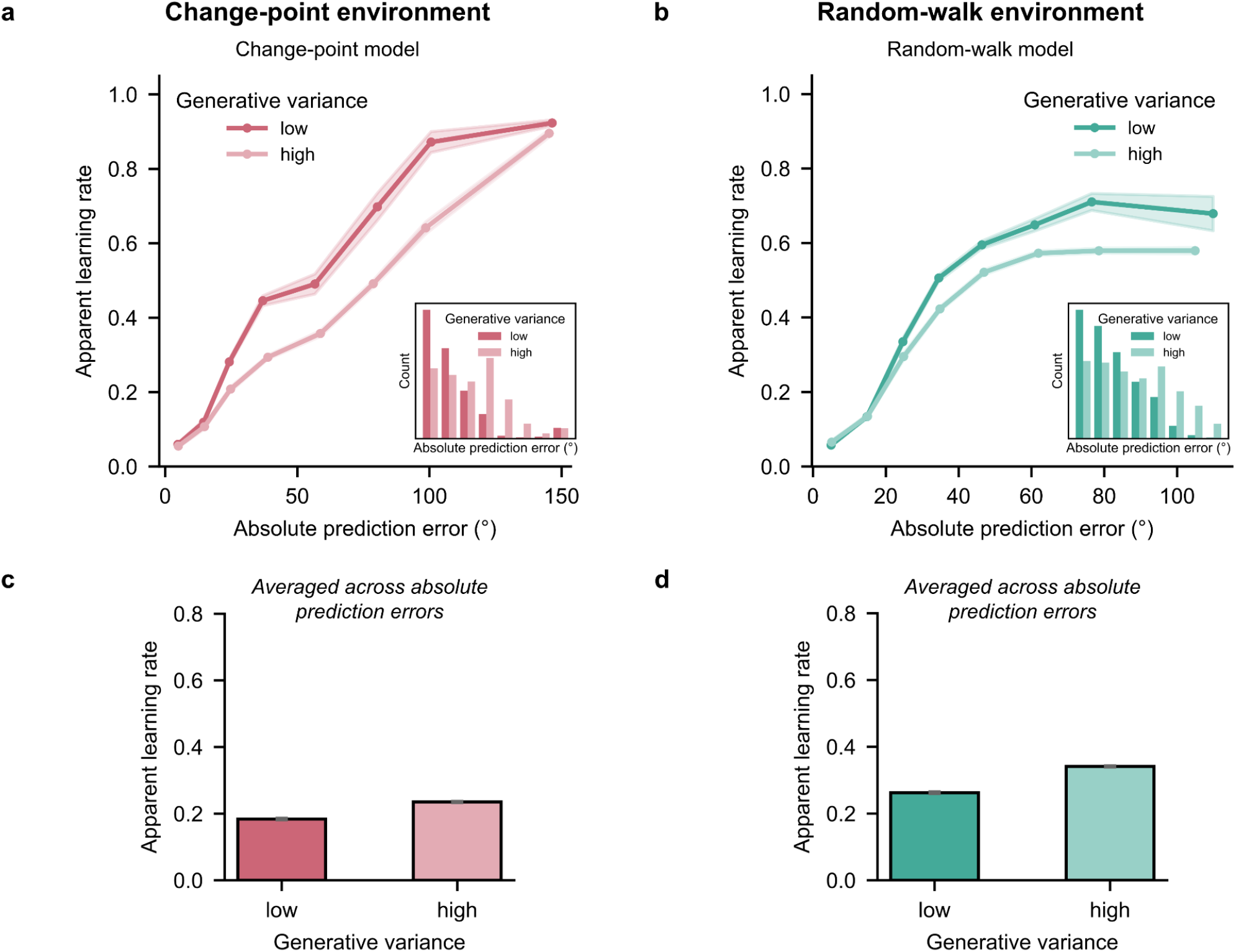
Model predictions: Environmental dynamics shape how apparent learning rates scale with prediction-error magnitude. (a, b) Apparent learning rate as a function of absolute prediction error for the change-point model in the change-point environment (a) and the random-walk model in the random-walk environment (b), split by generative variance. Insets show the corresponding prediction-error distributions. The change-point model predicts a sharper rise in apparent learning rate with prediction-error magnitude than the random-walk model. Both models show reduced apparent learning rates for small errors, reflecting lower movement probability. (c, d) Mean apparent learning rates under low and high generative variance for the same model–environment pairs. Although apparent learning rates are lower under high variance within matched prediction-error bins, their overall means are higher under high variance, owing to the shift toward larger prediction errors. Models were simulated on the experimental task sequences using one canonical response-parameter set shared across models and environments (see Methods). Shaded areas and error bars show ± s.e.m. across simulated participant sequences. See Supplementary Figure 1 for sensitivity to the response parameters and Supplementary Figure 2 for the equivalent simulations without the response model.

When prediction errors are very small (≤ 10°), both models show a large decrease in learning rate (Figures 3a and 3b). This suppression arises not from the inference process itself—beliefs continue to be updated—but from the response model. Because movement probability increases with prediction-error magnitude, the model rarely moves the paddle after minimal prediction errors. Therefore, many of these very-low-error trials produce no paddle movement, yielding a learning rate of zero, and lowering the average learning rate in this regime.

The effects of environmental stochasticity on apparent learning rate were more nuanced than might be anticipated from previous model predictions that learning rates should be lower under higher stochasticity (Piray & Daw, 2024; Pulcu & Browning, 2025). Within matched prediction-error bins, higher stochasticity reduced apparent learning rates in both models (Figures 3a and 3b), yet its net effect on average apparent learning rate was positive in both (Figures 3c and 3d). This positive overall effect arose in part because higher stochasticity generated larger prediction errors (insets in Figures 3a and 3b), which increased the learning rate calculated from target responses in the change-point model (Supplementary Figure 2) and movement probability in both models.

Environmental stochasticity therefore has multiple consequences for apparent learning rate, arising from both the inference component and the response model rather than from a single mechanism. Within the inference component, higher inferred stochasticity can lower the apparent learning rate calculated from target responses, but this effect interacts with other inferred quantities that also affect this measure, including change-point probability and volatility (Figure 3; Supplementary Figure 3). Beyond the target response, the response model can further modulate apparent learning rate because movement probability depends on prediction-error magnitude. Preceding response history introduces a further dependency: when the paddle is not moved across several trials, a later movement can express target changes accumulated over those trials. Higher stochasticity changes the distribution and sequence of prediction errors, which can affect apparent learning rate through both of these response-level dependencies. Modulation of apparent learning rate by environmental stochasticity therefore reflects the interaction of these processes and cannot be attributed to a single mechanism in the present models.

Having established the models’ distinct predictions, we next examined how human behavior compared with those predictions and, crucially, whether humans deployed the appropriate generative model for each environment. We addressed this question in two complementary ways: first, by fitting both models to each participant’s trial-by-trial responses and using cross-validation to determine which model better explained their behavior; and second, by testing whether human learning-rate adjustments mirrored the qualitative signatures predicted by the model matched to each environment.

### Model comparison shows that human behavior is best explained by the generative model matched to each environment

To compare the models formally, we fitted the change-point and random-walk models to each participant’s trial-by-trial responses and compared how well they predicted held-out responses (Figure 4; see Methods). We used 5-fold blockwise cross-validation: on each fold, the models were fitted on 12 of the 15 task blocks and evaluated using the log likelihood of responses in the remaining three blocks. Both models used the same stochastic response model, and only the response-model parameters were fitted; all parameters governing latent-state inference were fixed to the task-defined values. Differences in held-out likelihood therefore could not be attributed to unequal model flexibility, but instead reflected differences in how well the underlying inference models accounted for behavior. Group-level evidence was quantified with random-effects Bayesian model selection.

**Figure 4.**
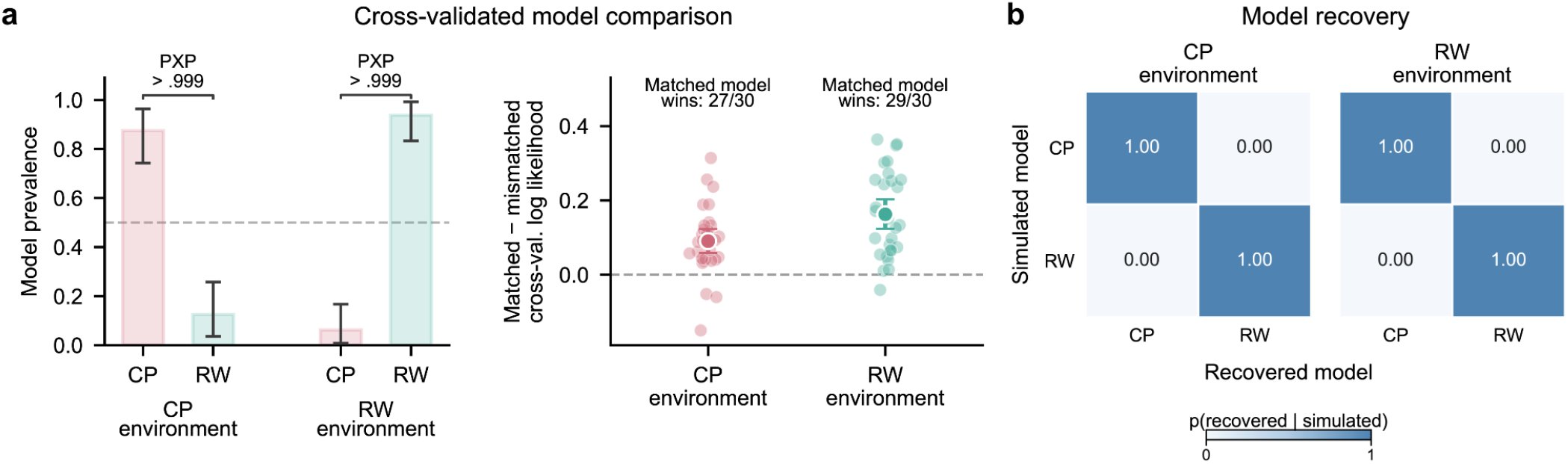
Human behavior is best explained by the model matched to each environment. (a) Cross-validated model comparison in the change-point (CP) and random-walk (RW) environments (n = 30 participants each). Left: model prevalence from random-effects Bayesian model selection. In each environment, the matched model had the higher prevalence (protected exceedance probability, PXP > .999). Right: difference in cross-validated log likelihood per response between the matched and mismatched models. The matched model better predicted held-out responses in nearly all participants (winner counts annotated). (b) Model recovery. The generating model was correctly identified in every simulated dataset (30 per generating model and environment), showing that the comparison reliably distinguishes the two models. In a (left), bars and error bars show posterior means and 95% credible intervals; in a (right), translucent circles show participants, and opaque circles and error bars show the mean and bootstrap 95% confidence interval. Dashed lines mark equal prevalence (0.5) and no difference (0). See Methods for the fitting, cross-validation, and model-recovery procedures, and Supplementary Figure 4 for parameter recovery.

The model matched to each environment best explained human behavior in nearly all participants (Figure 4a). The change-point model won in 27 of 30 participants in the change-point experiment (model prevalence 0.875, protected exceedance probability > 0.999), whereas the random-walk model won in 29 of 30 participants in the random-walk experiment (model prevalence 0.938, PXP > 0.999).

Model recovery and parameter recovery further validated the fitting and comparison procedure. The generating model was correctly identified for all simulated datasets (Figure 4b), and the four fitted response-model parameters were accurately recovered in both models (all Pearson correlations ≥ 0.99; see Supplementary Figure 4, which also shows the distributions of parameter estimates from full-data refits).

The formal comparison therefore showed that human behavior was better explained by the environment-matched model. We next examined whether participants’ learning-rate adjustments also mirrored its qualitative predictions.

### Human learning-rate adjustments mirror the qualitative signatures predicted by the environment-matched models

The models predicted that apparent learning rates should depend differently on prediction-error magnitude under the two environmental dynamics: specifically, they should increase more sharply in the change-point than in the random-walk environment (Figure 3). Human behavior showed the same dynamics-specific signature: apparent learning rates increased more sharply with prediction-error magnitude in the change-point environment than in the random-walk environment (Figure 5). To quantify this difference, we fitted a line to each participant’s learning rates across six common prediction-error bins spanning 10–110°. The mean participant-level slope per 10° was 0.064 ± 0.004 in the change-point environment and 0.033 ± 0.006 in the random-walk environment (difference = 0.030, 95% CI [0.017, 0.044], Welch’s t(50.8) = 4.56, p = 3.2 × 10⁻⁵, Cohen’s d = 1.18); over the same range, mean learning rates increased by 0.559 from the 10–20° bin to the 90–110° bin under change-point dynamics and by 0.279 under random-walk dynamics.

**Figure 5.**
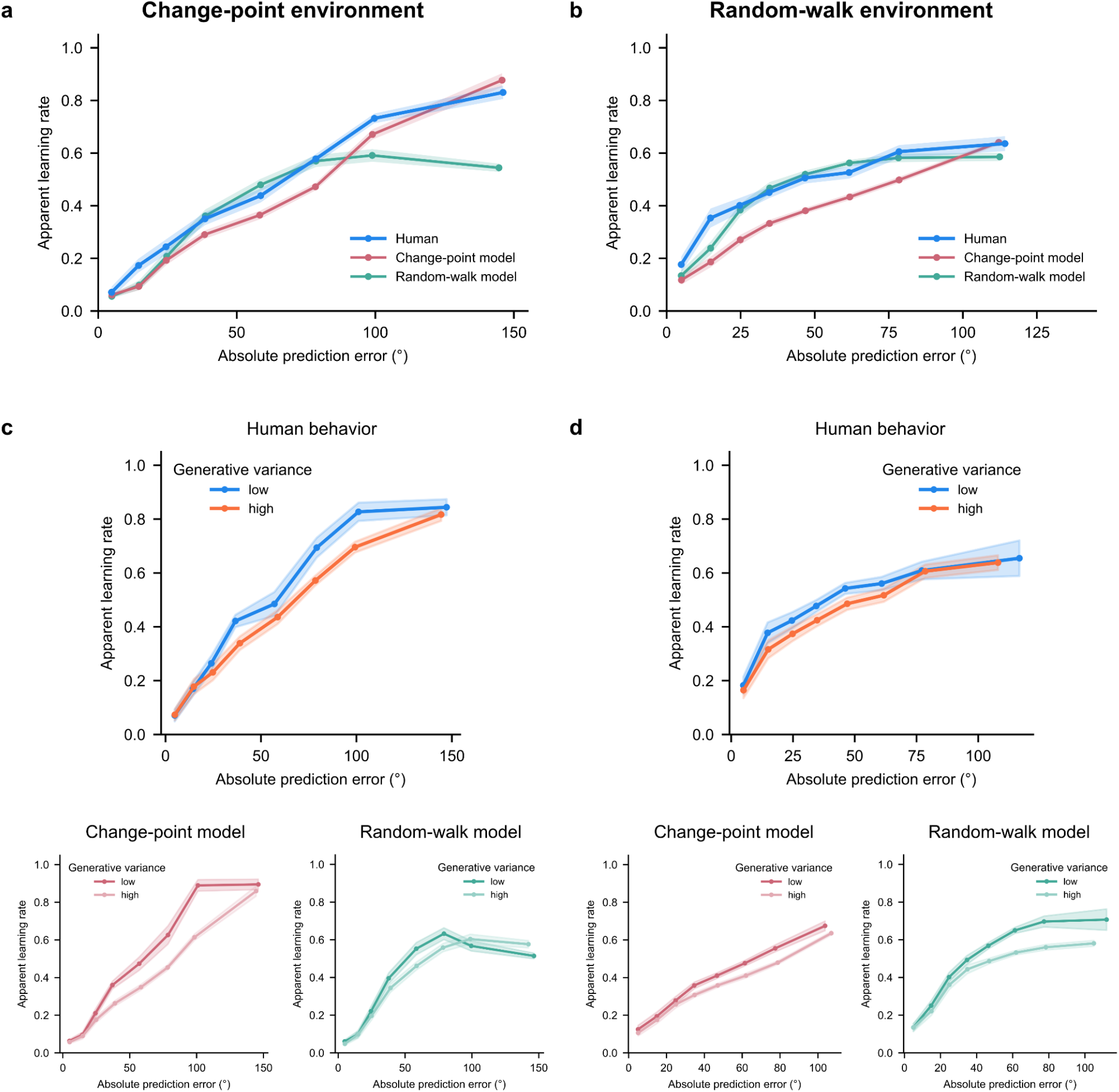
Human apparent learning rates follow the model matched to each environment. (a, b) Apparent learning rate as a function of absolute prediction error in the change-point (a) and random-walk (b) environments, overlaid with posterior-predictive simulations of the fitted change-point and random-walk models. Human apparent learning rate rises more steeply with prediction-error magnitude in the change-point than in the random-walk environment, and in each environment the human curve is closer to the matched than to the mismatched model (statistics in the main text). (c, d) The same data split by generative variance, for humans (top) and each fitted model (bottom). In both environments, human apparent learning rates are lower under high than low variance at matched prediction errors. Model curves were simulated on each participant’s task sequences using that participant’s fitted response parameters under the model shown (see Methods). Shaded areas show ± s.e.m. across participants.

We next asked whether these human learning-rate adjustments could be fully accounted for by response-parameter adaptation, or whether they also reflected environment-matched inference. Response-parameter adaptation can alter apparent learning-rate curves—for example, through a higher probability of initiating movements for small errors in the random-walk environment—without itself indicating different inference computations. Both fitted candidate models could accommodate such response differences because they used the same four-parameter response model, whose parameters were fitted separately for each participant under each candidate. However, they differed in their inference component. We therefore compared how closely each candidate reproduced each participant’s learning-rate curve in posterior-predictive simulations.

Human curves were closer to the matched than to the mismatched model in both environments (Figures 5a and 5b): the mean absolute error between the human curves and the matched model curve was 0.082 versus 0.107 with the mismatched model curve in the change-point environment (paired t(29) = 2.47, p = .020, Cohen’s d_z_ = 0.45) and 0.096 versus 0.125 in the random-walk environment (paired t(29) = 2.98, p = .006, d_z_ = 0.54). The conclusion was unchanged using mean squared error (change-point: p = .035; random-walk: p = .006).

Moreover, a consistent correspondence with model predictions was revealed after decomposing apparent learning rate into two components: (i) the proportion of trials on which participants moved their paddle (Supplementary Figure 5), and (ii) learning rate conditional on movement (Supplementary Figure 6; see also Supplementary Figure 7 for apparent-learning-rate distributions). At small prediction errors (≤ 30°), movement probability was 0.259 higher in the random-walk than in the change-point environment (Welch’s t(54.1) = 4.12, p < .001, d = 1.07). Both fitted candidates captured this response-level difference through their separately estimated response parameters.

In addition, the closer correspondence with the environment-matched model remained when learning rates were calculated only on trials in which participants moved their paddle (Supplementary Figure 6). Movement-conditional curves were closer to the matched than to the mismatched model in both the change-point environment (mean absolute error: 0.128 versus 0.169; paired t(29) = 4.66, p < .001, d_z_ = 0.85) and the random-walk environment (0.128 versus 0.155; t(29) = 2.40, p = .023, d_z_ = 0.44). The same conclusion was obtained using mean squared error (both p ≤ .008). Thus, the closer correspondence with the environment-matched model was not solely due to between-environment differences in how often participants moved their paddle.

Finally, we tested whether a Rescorla–Wagner learner augmented with the stochastic response model could account for these patterns. Although this model generated rising apparent learning rates from a constant latent rate, human curves were closer to the environment-matched Bayesian model for both raw and movement-conditional learning rates in both environments (paired comparisons: all MAE p ≤ .028; corresponding MSE p ≤ .013; Supplementary Figure 8). Consistent with this curve-level evidence, a recovery-validated, five-fold comparison with matched parameter flexibility favored the Bayesian model for 27/30 participants under change-point dynamics (model prevalence = 0.873, PXP > 0.999) and 20/30 under random-walk dynamics (prevalence = 0.662, PXP = 0.755). Thus, rising apparent learning rates do not by themselves imply rising latent learning rates, but both curve correspondence and cross-validated model comparison favored the Bayesian model.

### Effects of environmental variance and the type of environmental change on human learning rates

Across matched prediction-error bins, human apparent learning rates were lower on average during high-than low-variance periods in both environments (Figures 5c and 5d; participant-level high-minus-low differences were calculated within each prediction-error bin and then averaged across bins: change point, mean difference = −0.054, t(29) = −4.27, p < .001, d = −0.78; random walk, mean difference = −0.040, t(29) = −3.64, p = .001, d = −0.67). As the model analyses above show, however, this contrast does not by itself distinguish the inference and response processes that produced it.

Human apparent learning rates were also sensitive to the type of environmental change (Supplementary Figure 9). For each participant, the event-related effect was calculated as the apparent learning rate at the first observation following an event minus the mean apparent learning rate across the two preceding observations. In the change-point environment, the effect following a change point in the mean exceeded the much smaller effect following a variance increase by 0.465 ± 0.033 (paired t(29) = 13.98, p < 10⁻¹³, d_z_ = 2.55). The human response profile following mean changes was also closer to the fitted change-point model than to the fitted random-walk model (participant-level MSE across the first eight post-change observations: 0.009 versus 0.022, paired t(29) = 3.78, p < .001, d_z_ = 0.69; MAE gave the same conclusion, p = .002).

### Paddle-width adjustments reflect variance inference and distinguish variance changes from mean change points

We now turn to humans’ ability to track the current generative variance. This ability can be examined through participants’ paddle-width choices, which provide a behavioral measure of their variance estimates.

Humans’ paddle-width choices appropriately tracked the current generative variance in both environments: they more often chose the large paddle when variance was high and the small paddle when variance was low (Experiment 1, change-point environment: Figure 6c; average proportion of trials using the large paddle, mean ± s.e.m.: 17.9 ± 2.4% vs. 48.8 ± 3.9% in low- vs. high-variance periods, paired-samples t test: t(29) = −11.0, p = 7.0 × 10⁻¹²; Experiment 2, random-walk environment: Figure 6f; 37.9 ± 4.7% vs. 55.5 ± 5.1%, t(29) = −7.14, p = 7.4 × 10⁻⁸). The fitted environment-matched variance-inference models showed the same directional adjustment in both environments (Figures 6b and 6e), whereas the fitted fixed-low- and fixed-high-variance controls predominantly selected the paddle width favored by their fixed variance assumption (Figures 6a and 6d). In both environments, participants’ variance-dependent paddle-width adjustments were closer to the fitted environment-matched variance-inference model than to the fixed-variance controls. The absolute difference between human and model adjustments was smaller for the matched variance-inference model than for the fixed-low- and fixed-high-variance control models by 0.217 and 0.143, respectively, in the change-point environment (both t(29) ≥ 6.30, Holm-adjusted p ≤ 7.03 × 10⁻⁷, d_z_ ≥ 1.15), and by 0.106 and 0.087 in the random-walk environment (both t(29) ≥ 3.90, Holm-adjusted p < .001, d_z_ ≥ 0.71). Squared-difference and Wilcoxon comparisons gave the same conclusion.

**Figure 6.**
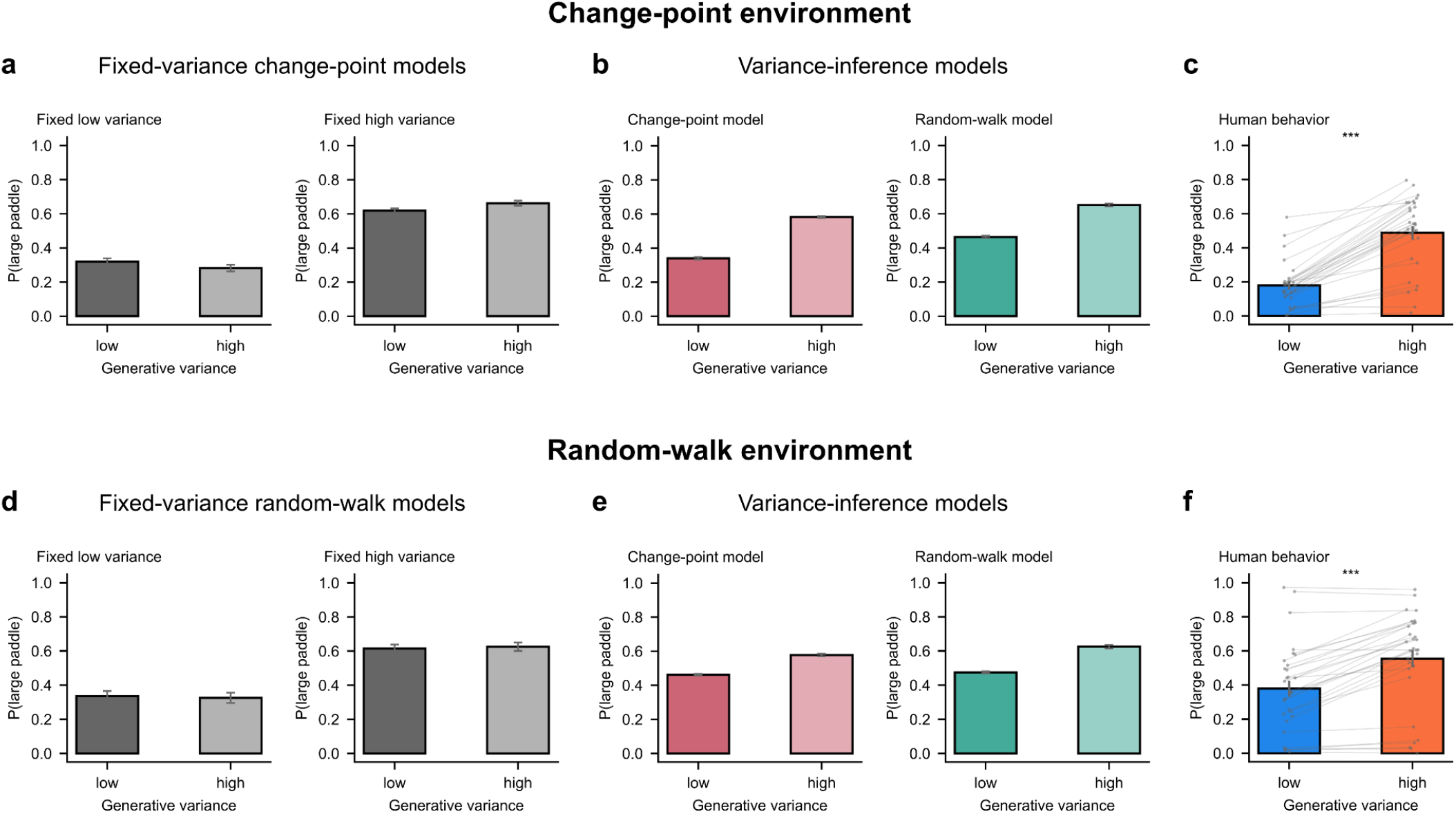
Human paddle-width adjustments reflect their inference of the current generative variance. Proportion of trials in which the large paddle was selected under low versus high generative variance, in the change-point (a–c) and random-walk (d–f) environments. (a, d) Fitted fixed-variance control models, which assume a fixed low or high variance, mostly select the paddle width favored by that assumption and barely adjust to the current variance. (b, e) Fitted change-point and random-walk models with variance inference select the large paddle more often when variance is high. (c, f) Humans show the same adjustment in both environments, and their adjustments are closer to the environment-matched variance-inference model than to the fixed-variance controls (statistics in the main text). Model bars were simulated on each participant’s task sequences using that participant’s fitted response parameters under the model shown (see Methods). Bars and error bars show means ± s.e.m. across participants; gray dots and lines show individual participants. ***p < .001, paired t-test comparing low versus high variance.

Participants’ paddle-width adjustments further showed that variance tracking occurred online within a sequence (Figure 7d). Their probability of selecting the large paddle increased progressively after variance increases and decreased progressively after variance decreases; the two curves crossed about three observations after the event. The fitted change-point model showed the same directional and sustained adjustments (Figure 7c).

**Figure 7.**
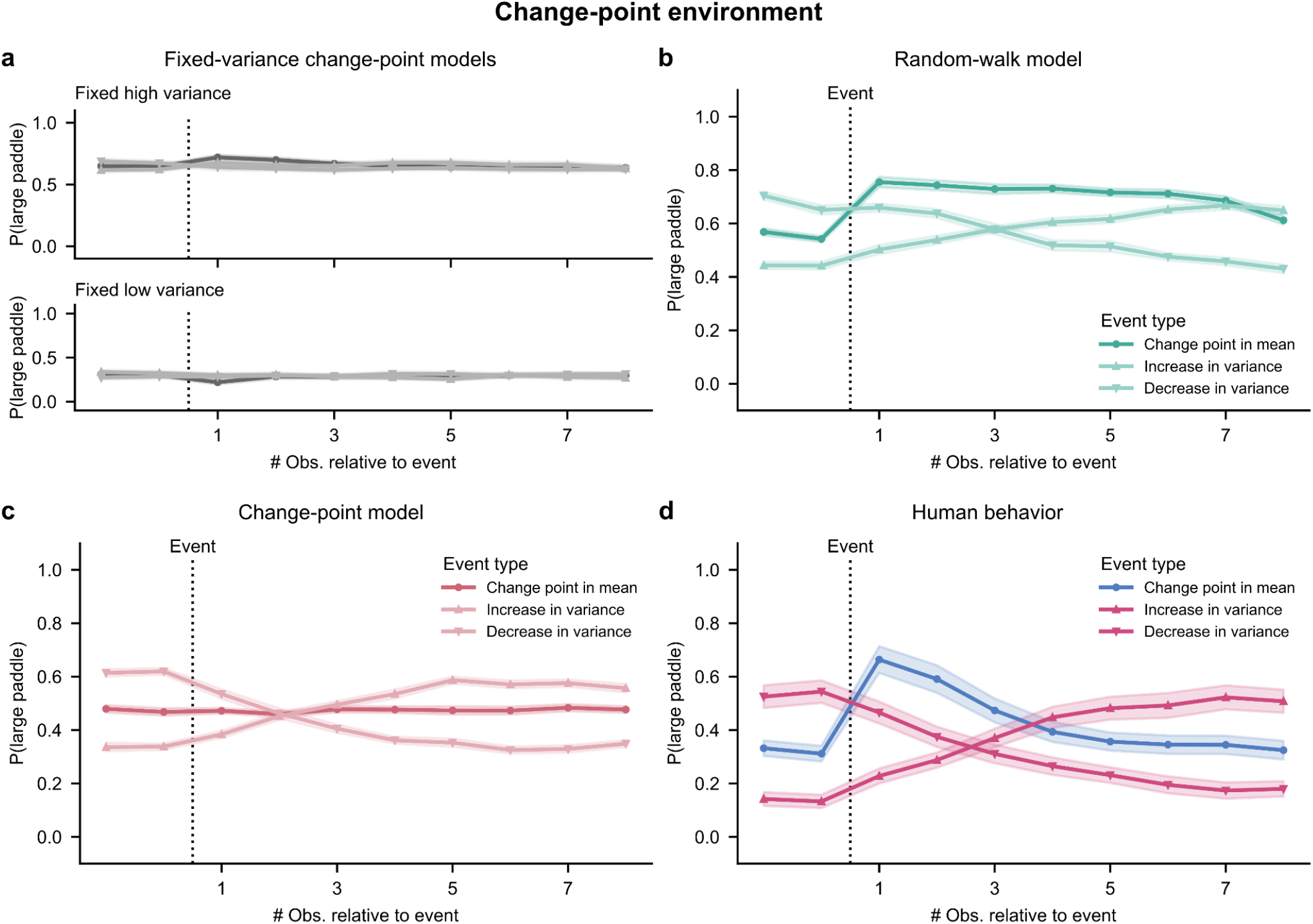
Human paddle-width choices distinguish changes in variance from change points in the mean. Proportion of trials in which the large paddle was selected around change points in the mean, increases in variance, and decreases in variance in the change-point environment. (a) Fitted fixed-variance change-point models remain near the paddle width favored by their fixed variance assumption after all three event types. (b) The fitted random-walk model adjusts its paddle width after variance changes, but also widens after mean change points and remains wide throughout the plotted window. (c) The fitted change-point model adjusts its paddle width after variance changes but not after mean change points. (d) Humans show sustained adjustments after variance changes but only a transient widening after mean change points (statistics in the main text). Model curves were simulated on each participant’s task sequences using that participant’s fitted response parameters under the model shown (see Methods). Dotted lines mark the event; shaded areas show ± s.e.m. across participants.

A key question was whether participants distinguished changes in variance from change points in the mean. A mean change point and an increase in variance can both initially produce observations that deviate substantially from the participant’s current mean estimate, making their sources difficult to distinguish from single observations. Previous work has suggested that humans may misattribute high outcome noise (high generative variance, in our terminology) to volatility, treating random variability as evidence that the mean is changing (Pulcu & Browning, 2025). We tested this distinction in the change-point environment, where both the mean and variance changed abruptly, allowing their effects on paddle width to be compared within the same task (Figure 7).

Human paddle-width responses distinguished changes in variance from changes in mean in the change-point environment (Figure 7d). Following an increase in variance, participants showed a sustained increase in large-paddle use, whereas a change point in the mean produced only a transient widening response. Among 27 of the 30 participants for whom both response durations could be estimated (in the remaining three, the post-event peak did not exceed baseline for at least one event type), widening after variance increases lasted longer than widening after mean change points (9.56 ± 0.56 vs. 2.26 ± 0.17 observations, mean ± s.e.m.; Wilcoxon signed-rank test: W = 0, p = 5.3 × 10⁻⁶; all 27 participants showed a longer duration after variance increases than after mean change points; 95% CI for this proportion [87.5%, 100%], binomial test p = 1.5 × 10⁻⁸; see Methods). This human profile contrasted with the fitted fixed-variance controls, whose paddle choices remained centered on the widths favored by their respective variance assumptions and showed no sustained direction-specific response to variance changes (Figure 7a). It also differed from the fitted random-walk model, which widened after a mean change point and remained elevated throughout the plotted post-event window instead of returning toward baseline after a brief response (Figure 7b). The fitted change-point model captured the dissociation between sustained responses to variance changes and the absence of sustained widening after mean changes, but did not capture the transient human widening (Figure 7c).

Finally, to test whether participants’ paddle-width choices could be explained by a single fitted subjective variance or were better explained by online inference of changes between two variance levels, we compared the environment-matched variance-inference model with a corresponding constant-variance variant based on their held-out paddle-width likelihoods (Supplementary Figure 10). The variance-inference model was strongly favored in the change-point experiment (26/30 participants; prevalence 0.859, PXP > 0.999; mean inferred-minus-constant cross-validated log-likelihood difference per response +0.047, bootstrap 95% CI [0.031, 0.063]). Evidence was more heterogeneous in the random-walk experiment: the variance-inference model was favored for 21/30 participants (prevalence 0.680, PXP = 0.801), but the participant-level cross-validated log-likelihood differences formed a visibly non-unimodal distribution, with six participants showing strongly negative values clearly separated from the remaining 24 (mean difference per response −0.034, bootstrap 95% CI [−0.126, 0.049]). This formal comparison therefore provides strong converging evidence for online variance inference in the change-point experiment and more modest evidence in the random-walk experiment.

### Hybrid model weights show a shift toward the model matched to each environment

To extend our results, we asked whether the environment-dependent shift toward the matched model would remain when the model space was expanded to allow the change-point and random-walk predictions to mix. We therefore fitted a hybrid model that combines the two models’ predictive distributions with a participant-specific weight *w* (*w* = 1 for the change-point component and *w* = 0 for the random-walk component; see Methods). The fitted weight provided a graded summary of how strongly each participant’s responses expressed the predictions of the two models.

The weight distributions shifted strongly between environments (Supplementary Figure 11a). The mean ± s.e.m. weight was 0.635 ± 0.042 (median = 0.637) in the change-point environment and 0.118 ± 0.025 (median = 0.079) in the random-walk environment (Mann–Whitney test: *U* = 874.0, p = 3.0 × 10⁻¹⁰; rank-biserial correlation = 0.942). Thus, allowing the models’ predictions to mix preserved the strong environment-dependent difference found in the earlier model comparison.

To investigate which components of the response contributed to this environment-dependent shift, we refitted the hybrid model separately to paddle-location and paddle-width responses (Supplementary Figure 11b–c). The results reflected the models’ structural difference and the preceding behavioral findings. Location-only weights differed strongly between the change-point and random-walk environments: the means ± s.e.m. were 0.660 ± 0.035 (median = 0.676) and 0.120 ± 0.030 (median = 0.010), respectively (*U* = 869.0, p = 4.1 × 10⁻¹⁰; rank-biserial correlation = 0.931). In contrast, width-only weights did not differ reliably: the means ± s.e.m. were 0.616 ± 0.048 (median = 0.559) and 0.547 ± 0.056 (median = 0.500), respectively (*U* = 528.0, p = 0.250; rank-biserial correlation = 0.173). The models differ in the dynamics of the latent mean, which is expressed primarily through paddle-location adjustments, but assume the same switching low- and high-variance states. Both models predict paddle widening following increases in variance and narrowing following decreases, but their paddle-width predictions diverge most clearly after a change point in the mean: the fitted random-walk model shows sustained widening, the fitted change-point model shows no sustained widening, and participants show the brief transient widening reported in Figure 7. The location-only weights are therefore consistent with the environment-dependent differences in learning-rate adjustments, whereas the width-only weights primarily reflect variance tracking common to both models; the brief human width response after a mean change lies between their predictions.

Note that these fitted weights do not establish that participants used a hybrid strategy. The hybrid model cannot distinguish a stable mixture of the two strategies from trial-to-trial variation in their use, and a less extreme weight may partly capture occasional deviations from one component model rather than consistent use of both strategies.

## Discussion

Humans constantly face environments that change in different ways—sometimes abruptly, sometimes gradually. In this study, we asked whether and how humans adapt their learning to such distinct forms of environmental dynamics. Cross-validated model comparison showed that participants’ trial-by-trial responses were best explained by the Bayesian model whose generative assumptions matched the environment (Figure 4). Consistent with this result, participants adjusted their learning rates in systematically different ways across change-point and random-walk environments, in close alignment with the predictions of the corresponding environment-matched fitted models (Figure 5). We further asked whether participants could distinguish different types of latent environmental changes—either in mean, or in variance. This was evident in their simultaneous adjustments of paddle location and paddle width, which jointly tracked the two latent variables; in the change-point environment, their width responses further distinguished changes in variance from changes in mean (Figures 6 and 7). Together, these results advance previous work by providing a direct comparison of change-point and random-walk dynamics within a unified framework, and showing that humans flexibly deploy world models suited to the environment’s dynamics and its changing latent properties.

Notably, the same prediction error elicited different adjustments under change-point versus random-walk structure. In the change-point environment, large prediction errors (PEs) triggered sharp, transient increases in apparent learning rate, consistent with detecting discrete changes in the latent mean. In contrast, in the random-walk environment, the function mapping from PEs to learning rates was flatter, with more even weighting of small and large PEs, reflecting steadier, continuous integration of new evidence appropriate for gradual drift. By directly contrasting these regimes within the same paradigm, the present findings extend and unify two largely independent literatures on adaptive learning under change points (e.g. (Meyniel et al., 2015; Nassar et al., 2010)) and random walks (e.g. (Mathys et al., 2011; Piray & Daw, 2024)). Critically, together with the formal model comparisons, these dynamics-specific patterns show that learners do not rely on a fixed, error-driven update rule, but instead condition their interpretation of prediction errors on the environment’s dynamics—a hallmark of world-model accounts of learning (Bruckner, Heekeren, et al., 2025).

An important contribution of our framework is that it separates latent inference from stochastic response generation. Modeling response generation explicitly allows response-level processes to be distinguished from underlying inference. This distinction is important when interpreting apparent learning rates calculated from overt responses because a prediction-error-dependent response process can cause apparent learning rate to vary with prediction-error magnitude even when the latent learning rate is constant (Supplementary Figure 8).

Our results help clarify apparent contradictions in the literature on learning by showing that stochasticity does not exert a unitary effect on apparent learning rate. Instead, its influence depends on the assumed dynamics and interacts with prediction error. Higher stochasticity produces more frequent large PEs, which can raise average apparent learning rate even though it lowers apparent learning rate when PE magnitude is held constant (Figure 3). Our computational analyses show why the size and direction of this effect depend on the assumed generative model and response process (Figure 3; Supplementary Figure 3). This account may help reconcile the opposite effects of stochasticity reported in prior studies (Piray & Daw, 2024; Pulcu & Browning, 2025). Differences in task instructions between the studies—particularly the extent to which participants were informed about the task’s underlying generative structure—may have contributed to these contrasting results. For example, uninstructed participants in the latter study may have treated large PEs as possible change points (Pulcu & Browning, 2025). This remains a hypothesis and may coexist with Pulcu and Browning’s stochasticity–volatility misattribution account.

Beyond inferring the mean, participants were able to track latent variance online in both environments and, in the change-point environment, dissociate changes in variance from changes in mean. This was evidenced by coordinated adjustments in paddle location and paddle width, which tracked changes in the latent mean and variance, respectively. Following variance change points, participants rapidly recalibrated their paddle width to the new variance—often within a few trials—and then maintained this calibration thereafter. By contrast, mean change points elicited only a brief uptick in paddle width and, critically, a step-like shift in the paddle location, manifested as a transient spike in the apparent learning rate—consistent with the fitted change-point model (Figure 7 and Supplementary Figure 9). These findings extend prior work on uncertainty and confidence during learning (e.g., (Boldt et al., 2019; Meyniel et al., 2015; Nassar et al., 2010; Vaghi et al., 2017)) and complement recent studies on stochasticity inference (e.g., (Piray & Daw, 2024; Pulcu & Browning, 2025)) by demonstrating within-block, online adaptation.

### Human behavior in our task reflects model-based learning through probabilistic inference

World-model accounts of learning argue that learners use an internal generative model of the environment to interpret prediction errors and guide belief updates (Bruckner, Heekeren, et al., 2025). Prediction errors are not treated as uniform learning signals, as in model-free algorithms, but are instead evaluated in light of the learner’s internal generative model of the environment’s dynamics. This context-dependent interpretation of error explains the dynamics-specific learning-rate patterns described above, and situates our findings within broader debates contrasting model-free and model-based mechanisms of learning (Collins & Cockburn, 2020; Daw et al., 2005; Dolan & Dayan, 2013; Doody et al., 2022; Feher da Silva & Hare, 2020; Gläscher et al., 2010). While both forms of learning likely coexist in the brain, our results provide evidence for environment-sensitive, model-based inference. Moreover, participants were able to deploy the generative model matched to the environment immediately, without environment-specific training or feedback, by relying on abstract structural knowledge acquired through instructions. This capacity to apply an appropriate inference or learning strategy without prior task exposure connects to work on theory-based learning in humans (Lake et al., 2015; Tsividis et al., 2026) and demonstrations of zero-shot learning in artificial systems (Brown et al., 2020; Kojima et al., 2022).

The convergence between human and model predictions highlights the importance of aligning computational models with the environment’s generative structure. The environment-matched models best predicted participants’ held-out responses and reproduced the distinct learning patterns observed across environments, manifested in trial-by-trial adjustments in learning rate and reported uncertainty. This alignment shows that differences in observed learning behavior—often interpreted as changes in learning mechanisms (Eckstein et al., 2022)—can arise naturally from Bayesian inference under different world models. Models tied to one class of dynamics, such as the Kalman filter, the Hierarchical Gaussian Filter and its extensions (Diaconescu et al., 2014; Kalman, 1960; Mathys et al., 2011; Weber et al., 2023), or change-point models (Adams & MacKay, 2007; Nassar et al., 2010), capture important aspects of learning but cannot by themselves explain humans’ ability to flexibly adjust computations to match the environment’s dynamics, and risk misattributing behavior when model and environment dynamics are mismatched. Our framework instead offers a unified lens for capturing behavior across environments by explicitly modeling assumptions about environmental structure within a common probabilistic-inference account. Given that exact Bayesian inference is in general computationally intractable, an open question concerns how humans approximate and implement it (Lieder & Griffiths, 2019; Van Rooij, 2008). Future work could compare normative predictions with process-level approximations, such as small recurrent neural networks (Foucault & Meyniel, 2021; Ji-An et al., 2025), hybrid neural-cognitive models (Eckstein et al., 2026), or other resource-bounded models (Bhui et al., 2021; Bruckner, Heekeren, et al., 2025; Bruckner, Nassar, et al., 2025; Drevet et al., 2022; Findling et al., 2019; Franklin & Frank, 2015; Gershman et al., 2015; Murphy et al., 2021; Zhang et al., 2023), to uncover the mechanisms that enable human brains to approximate Bayesian inference under resource constraints. Process-level models could further characterize individual differences in the computations underlying probabilistic inference (Mikus et al., 2025; Piray & Daw, 2021; Weber et al., 2023).

The stochastic response model captures a non-normative feature of participants’ behavior separately from latent inference: they often left the paddle unchanged, especially after small prediction errors. Such perseveration may reflect a cost associated with initiating or executing a response, a satisficing or resource-rational policy in which a new response is generated only when the current one is no longer “good enough,” or a combination of these factors (Bruckner, Nassar, et al., 2025; Gershman, 2020; Nassar et al., 2021; Simon, 1956). In a related task in which the starting position of each response was randomly displaced from the previous response, participants rarely repeated their previous response or left the response at its displaced starting position, but their responses were biased toward that position (Bruckner, Nassar, et al., 2025). The authors attributed this pattern to resource-rational sampling rather than movement costs. Our task did not include this manipulation, so it does not distinguish between these accounts. How responses are made in a given task may also influence the relative contributions of these factors.

A future direction that remains to be investigated is how humans infer environmental structure, including its dynamics, when this is not scaffolded by instruction. In our study, participants were explicitly informed about the task’s generative structure, as our study was focused on estimating latent states (namely, the generative mean and variance) rather than on discovering which structural form governed the environment. The present findings should therefore be interpreted within this instructed-context boundary condition. An open question is whether the dynamics-specific learning signatures observed here re-emerge once learners have discovered the underlying structure. This broader problem, known as *structure learning* (Gershman & Niv, 2010), involves identifying which generative process best explains incoming observations and when to switch between alternative world models, and likely engages additional inference mechanisms beyond those studied here. Recent work has begun to reveal such structure-learning mechanisms (Gershman et al., 2017; Tomov et al., 2018, 2023). Extending the present framework to situations where environmental dynamics must themselves be inferred would help unify adaptive learning and structure discovery under a common computational account.

Two further extensions follow naturally from our design. First, a within-participant version of the task, with each participant experiencing both dynamics in counterbalanced order, would allow adaptation across the two dynamics to be tracked within individuals. Second, complementing paddle width with explicit confidence reports and response times could provide additional, potentially more graded indices of uncertainty and help test whether subjective uncertainty predicts the trial-by-trial learning-rate adjustments observed here (Boldt et al., 2019; Meyniel et al., 2015; Vaghi et al., 2017).

### Implications and outlook

Our findings establish a computational account of how environmental dynamics shape human learning. Theoretically, they demonstrate that humans flexibly deploy computational strategies tailored to the structure of their environment, providing strong evidence for adaptive, model-based cognition. Methodologically, they underscore the importance of aligning computational models with the generative structure assumed or inferred by the learner and illustrate how our framework provides a general approach for unifying analyses across different forms of environmental change. In addition, they generate concrete neural predictions for future work. Specifically, changes in mean versus variance should produce dissociable neural signatures, reflecting distinct forms of latent-state updating. Moreover, change-point environments should evoke transient, large neural responses to surprising events (e.g., pupil dilation, P3 amplitudes, and prefrontal and anterior cingulate cortex activity), whereas random-walk environments should elicit steadier, graded error-related responses. While prior work has investigated these neural signatures separately (Chakroun et al., 2020; Daw et al., 2006; Findling et al., 2019; Jepma et al., 2016; Kao et al., 2020; McGuire et al., 2014; Murphy et al., 2021; Nassar et al., 2012; O’Reilly et al., 2013), neural responses to change points versus random walks have not yet been compared directly.

## Methods

### Participants

Participants were recruited via Prolific, an online participant recruitment platform for academic research studies, and completed the task online in their web browsers. Data were collected in September 2024 for Experiment 1 and April 2025 for Experiment 2. The study was approved by the University of Oxford’s Central University Research Ethics Committee (CUREC Approval Reference: R96268/RE001). All participants provided informed consent prior to participation.

A total of 60 participants completed the study (30 in each of the two experiments), with a mean age of 35.0 years (SD = 11.9); fourteen identified as female and one did not report their gender. All participants reported normal or corrected-to-normal vision and fluency in English. All participants completed a comprehension quiz (repeating instructions if necessary until passing) and went on to complete all 15 blocks of the behavioral task. Participants received a base payment of £8.40 plus a performance-based bonus of up to £4.20. No participants were excluded from analyses.

### Task and experimental design

#### Task instructions

Participants completed a behavioral task presented as a “capture-the-beams” space exploration game. Before starting, they received instructions explaining the task through a cover story, describing the task’s goal, mechanics, and underlying generative process. Their objective was to maximize points, which in the cover story corresponded to energy collected for their colony, awarded for capturing beams emitted from a star.

The key concepts of *generative mean* and *variance* were introduced using simple analogies (a flashlight emitting beams with a certain average direction and spread). The two variance levels were described as two distinct states of the star (a focused state vs. a diffuse state). Participants were then introduced to the paddle and its two possible sizes, along with the point system. They had the opportunity to practice moving the paddle and switching sizes.

The amount of points awarded per observed beam depended on capture outcome and paddle size (+10 points for capturing with the small paddle, +5 points for capturing with the large paddle, -10 points for a miss). This scoring scheme created a trade-off between the higher likelihood of catching a beam with the large paddle and the greater reward per beam captured with the small paddle. Normatively, this incentivized participants to use the large paddle when the generative variance was high and the small paddle when variance was low.

The *dynamics* of the generative mean and variance were then explained. In Experiment 1, participants were told that the star’s average direction could change abruptly from time to time; in Experiment 2, they were told it changed gradually over time. In both experiments, they were told that the star’s state could change occasionally (switching between the focused, low-variance state and the diffuse, high-variance state). Participants were not informed about the frequency or magnitude of changes, nor about the precise variance levels.

At the end of the instructions, participants completed a short comprehension quiz that tested their understanding of how beam direction and spread changed and how paddle size influenced scoring. Those who did not pass the quiz reviewed the instructions until they succeeded, after which they proceeded to the task.

#### Task structure

Each participant completed 15 task blocks of 100 trials each. On each trial, participants could freely adjust the angular location of the paddle on a circular ring (0–360°) and select between the small paddle (60° wide) or the large paddle (120° wide). The paddle could be controlled using either the keyboard (left/right arrows to rotate, up/down arrows to change paddle size) or the mouse or touchpad (cursor movement for rotation, and scroll to change size). Once satisfied, participants confirmed their choice using the space bar or a mouse click.

After participants’ confirmation, the beam was revealed, and points were awarded according to whether it was captured and with which paddle size (as per the +10/+5/-10 scheme described above). Visual feedback showed whether the beam was captured or missed and how many points were gained or lost. The next trial then began.

At the start of each block, participants began with an initial score of 110 points. This starting score incentivized them to avoid losses from the outset, since points translated to a monetary bonus that could only be positive. At the end of each block, participants were shown their total score for that block and could take a break if they wished. Once they confirmed (by pressing the space bar), the next block began.

Participants were free to take as much time as they wished to make their choices. On average, they spent 2 min 30 s (± 4 s) per block in Experiment 1, and 2 min 34 s (± 5 s) per block in Experiment 2, corresponding to approximately 1.5s per trial in both cases.

#### Generative process and parameters

The observation sequences (beam angles) presented in each block were generated using a stochastic process, identical across the two experiments except for the dynamics of the generative mean. On each trial, the current generative mean and variance were updated, and a new observation (beam angle) was drawn. The generative process is fully described as follows.

- **Observation.** Each observation was drawn from a wrapped Gaussian distribution with the current generative mean and variance parameters.
- **Dynamics of the generative mean**.

○ Experiment 1: The mean followed a change-point process, with probability p = 1/16 of a change point (jump to a new value, sampled uniformly from 0–360) occurring on each trial.
○ Experiment 2: The mean followed a wrapped Gaussian random walk with volatility (process noise variance) of 15^2^.
- **Dynamics of the generative variance.** The generative variance switched between two levels, low (10^2^) and high (30^2^), with probability p = 1/16 of a switch occurring on each trial.
- **Initialization.** The generative mean was initially sampled uniformly from 0–360, and the generative variance was initially set randomly to low or high with equal probability

To optimize experimental efficiency, we imposed additional constraints on the sequences used in the experiment. Run lengths (intervals between change points) were constrained to 5–40 trials (avoiding excessively long or short stable periods) and change points in mean were required to elicit a change of at least 90° (avoiding excessively subtle change points). These constraints were not disclosed to participants or computational models.

Parameters such as paddle widths, variance levels, and reward values were selected based on simulations of Bayesian ideal observers. Simulations verified that the normative model would use both paddle sizes meaningfully (favoring the large paddle in high-variance periods and the small paddle in low-variance periods) and that change points in mean rarely caused the normative model to change its paddle size. This ensured that paddle size choices primarily reflected variance inference rather than mean uncertainty.

#### Implementation

The task was implemented in JavaScript and ran in participants’ web browsers. Participants accessed the task through a dedicated URL. The task was hosted on Pavlovia servers, ensuring compliance with the EU General Data Protection Regulation (GDPR).

The full experiment code, which can be used to run and test the task, is available at our GitHub repository (see *Code and data availability* for access details).

### Computational Modeling

We developed a family of Bayesian models. All models share the same four-stage architecture but differ in their assumptions about the generative process. Additional generative-model structures and parameterizations can be explored using our online code (see *Code and data availability* for access).

#### Model variants

The main Bayesian model set crosses two assumptions about mean dynamics (abrupt change points versus gradual random walks) with two assumptions about the generative variance (switching and inferred online versus fixed). The fixed-variance branch comprises separate low- and high-variance variants. This gives four model families:

- **Change-point model.** The environment-matched inference model for Experiment 1 assumes the change-point mean dynamics and switching variance described in *Generative process and parameters*.
- **Random-walk model.** The environment-matched inference model for Experiment 2 assumes the random-walk mean dynamics and switching variance described there, and additionally infers the volatility of the mean random walk rather than treating it as known.
- **Fixed-variance change-point model.** This family assumes the same change-point mean dynamics as the change-point model, but holds the generative variance fixed. The fixed-low- and fixed-high-variance variants assume a generative SD of 10° and 30°, respectively.
- **Fixed-variance random-walk model.** This family assumes the same random-walk mean dynamics as the random-walk model, but holds the generative variance fixed. The fixed-low- and fixed-high-variance variants assume a generative SD of 10° and 30°, respectively.

The Rescorla–Wagner and hybrid models are described below as control and extension models.

#### Model computations

On each trial, each model generates a response, *y_t_* , from the current observation, *x_t_*, through the four-stage architecture shown in Figure 2a. First, Bayesian inference updates the posterior over the latent state under the model’s assumed generative process. Second, the model uses this posterior to obtain the predictive distribution over the next beam location. Third, the model selects the target response, *y_t_*\*, comprising the paddle location and width that maximize expected reward under the predictive distribution. Finally, the stochastic response model generates the actual response from this target through a movement-initiation decision—whether to stay at the previous paddle location or move—followed by generation of the paddle location and width conditioned on this decision.

#### Mathematical details

##### Latent-state space

Each model’s latent state consists of the variables in its assumed generative process that are not directly observed. The change-point model represents *s_t_* = (µ*_t_*, σ*_t_*), where µ*_t_* ∈ [0°, 360°) is the generative mean and σ*_t_* ∈ {10°, 30°} the generative SD. The random-walk model represents *s_t_* = (µ*_t_*, σ*_t_* , ν*_t_*), additionally treating the random-walk SD (volatility) as an unknown latent variable, ν*_t_* ∈ {10°, 30°}. The fixed-variance controls instead represent only µ*_t_* and hold σ fixed at 10° or 30°, respectively.

##### Latent-state dynamics and observation model

The state-transition distribution *p*(*s_t+1_* | *s_t_*) and the observation distribution *p*(*x_t_* | *s_t_* ) use the corresponding dynamics and wrapped-normal observation process specified in *Generative process and parameters* above. That is, each model assumes the mean dynamics associated with its variant (change-point or random-walk) and, where applicable, the switching low- and high-variance states. The random-walk model also treats ν*_t_* as a switching latent variable with the same two-state dynamics as σ*_t_*.

##### Bayesian inference

Each model uses Bayesian filtering to infer the latent state of its assumed generative process from the observed beam locations (Foucault, 2023; Särkkä & Svensson, 2023):

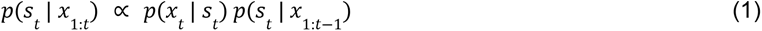

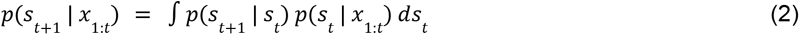

Here, *s_t_* is the latent state at time *t* and *x*_1:*t*_ is the sequence of observations up to that time. Equation (1) updates the posterior after the current observation, and equation (2) uses that posterior to obtain the predictive distribution over the next latent state. These recursive updates use the current observation and the posterior from the preceding trial, enabling online inference without retaining the full observation history.

To make filtering tractable, the circular mean μ was discretized on a 1° grid (360 values), while *σ* and *ν* each took the two discrete candidate values. Within this finite state space, filtering was exact: the posterior update was computed at every grid point, the integral in equation (2) was evaluated as a finite sum, and the complete joint distribution over the latent state was represented explicitly at each step (e.g., 360 × 2 values for the change-point model). No parametric, moment-matching, or reduced approximations, such as those used in approximate Bayesian delta-rule models (Nassar et al., 2010), were applied.

##### Prediction

The predictive distribution over the next observation is:

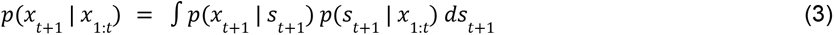

The two factors in the integrand are the observation distribution *p*(*s_t_*_+1_ | *x_t_*_+1_) and the posterior distribution over the next latent state *p*(*s_t_*_+1_ | *x*_1:*t*_) (equation (2)).

##### Target response

Candidate responses comprise every combination of a paddle location (discretized on a 0.1° grid) and one of the two available paddle widths. For each candidate, the model computes the expected reward under the predictive distribution and selects the candidate with the highest expected reward as the target response:

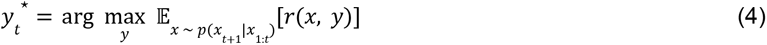

Here, *r*(*x*, *y*) is the task reward function. We denote the target response by *y_t_*\* = (*a_t_* *, *w_t_*\* ) and the actual response by *y_t_* = (*a_t_* , *w_t_* ), where *a* and *w* denote paddle location and width, respectively.

##### Stochastic response model

The stochastic response model generates *y_t_* from *y_t_*\* and has the same form for all inference models. On each trial, it first determines whether to stay at the previous paddle location or move. Let *angdiff(u, v)* denote the signed shortest circular difference from *v* to *u*, in the range [−180°, 180°). The signed prediction error, δ*_t_*, and its absolute magnitude, *d_t_* , are:

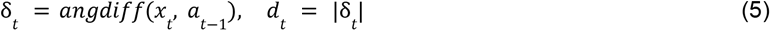

Here, *a_t_*_−1_ is the previous paddle location, before observation *x_t_*.

Movement is initiated with probability:

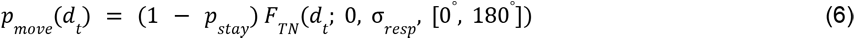

where *F_TN_* is a truncated-normal CDF, *p_stay_* is the stay-lapse probability (which sets the upper asymptote of *p_move_* to 1 − *p_stay_*), and σ*_resp_* is the movement-initiation scale in degrees. Larger values of σ*_resp_* reduce the probability of initiating movement at a given prediction error.

Conditioned on the movement-initiation decision, the paddle location and width are generated as follows. If movement is initiated, the paddle location follows a von Mises distribution centered on the target location (concentration κ*_move_*, parameterized as a circular SD in degrees σ*_move_*), and the paddle width equals the target width with probability 1 − ɛ*_w_*, where ɛ*_w_* is the width-choice error probability. If movement is not initiated, the paddle location remains at its previous value (von Mises with concentration fixed to 3283, i.e. with circular SD σ*_stay_* fixed to approximately 1°, absorbing recording imprecision around an unchanged location), and the width equals the previous width with probability 1 − ɛ*_w_*.

The response model has four fitted parameters: stay-lapse probability *p_stay_*, movement-initiation scale σ*_resp_*, movement circular SD σ*_move_*, and width-choice error probability ɛ*_w_*. Their estimation is described in *Model fitting and comparison* below.

Unless specified otherwise, model simulations shown in the paper used response-parameter values fitted separately for each participant under the model being simulated. Figure 3 and panels d–f of Supplementary Figure 3 used one canonical response-parameter set shared across both environments and models (*p_stay_* = 0.05, σ*_resp_* = 25°, σ*_move_* = 15°, ɛ*_w_* = 0.2). These values were chosen to be representative of the fitted parameter distributions: each lies within the interquartile range of the corresponding parameter across the pooled matched-model fits. In a supplementary one-at-a-time sensitivity analysis, we varied each parameter over a broader range informed by those distributions; Supplementary Figure 1 shows that the qualitative predictions remained consistent across these parameter settings.

##### Apparent learning rate

Apparent learning rate is not an internal variable of the Bayesian models. It was instead calculated from their simulated paddle responses using the same trial-wise measure applied to participant responses (see *Data analyses* below).

#### Model fitting and comparison

We fitted the candidate models to each participant’s trial-by-trial paddle responses to determine which inference model best accounted for behavior and to estimate participant-specific response parameters for subsequent simulations. We first estimated each model’s parameters from the response likelihood, then compared the models using blockwise cross-validation, and finally validated parameter estimation and model comparison through parameter recovery and model recovery, respectively (Wilson & Collins, 2019).

The likelihood was defined over the participant’s joint paddle-location and paddle-width responses. Let θ = (*p_stay_*, σ*_resp_*, σ*_move_* , ɛ*_w_*) denote the four response parameters. For participant *i* and candidate model *m*, the log likelihood for one block of *N* trials was:

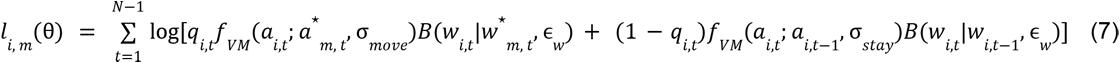

Here, *q_i,t_* = *p_move_*(*d_i,t_*) is the movement probability defined in equation (6), *f_VM_* is the von Mises density for paddle location, and *B*(*w*|*w*’, ɛ*_w_*) equals 1 − ɛ*_w_* when *w* = *w*’ and ɛ*_w_* otherwise. The likelihood is thus the sum of a movement term and a no-movement term, weighted by the movement probability and its complement, respectively. The movement term is centered on the candidate model’s target response (*a*\**_m,t_*,*w*\**_m,t_*), whereas the no-movement term is centered on the participant’s previous response (*a_i,t_*_-1_, *w_i,t_*_-1_). Log likelihoods were summed across the blocks included in each fit or evaluation.

We fitted the four response parameters θ = (*p_stay_*, σ*_resp_* , σ*_move_*, ɛ*_w_*) separately for each participant and model. We obtained maximum-likelihood estimates using SciPy’s ‘scipy.optimize.minimize’ function with five deterministic starting points spanning the parameter bounds. Powell’s method was used to optimize the negative log likelihood from each starting point, and the solution with the highest likelihood was retained. Optimization was performed in transformed space (log scales for positive parameters and logit scales for probabilities), with bounds *p_stay_* ∈ [0. 0001, 0. 59], σ*_resp_* ∈ [1^◦^ , 60^◦^ ], σ*_move_* ∈ [0. 5^◦^ , 40^◦^ ], and ɛ*_w_* ∈ [0. 0001, 0. 5]. Each optimization allowed up to 1,000 iterations; a non-converged solution was retried from its best parameter values with a limit of 3,000 iterations.

We compared the candidate models by 5-fold blockwise cross-validation. The 15 task blocks were partitioned into five folds of three blocks; for each fold, parameters were estimated from the remaining 12 blocks and equation (7) was evaluated on the held-out blocks. The folds were identical across models, and the five held-out log likelihoods were summed for each participant and model. Group-level evidence was quantified by random-effects Bayesian model selection applied to these summed cross-validated log likelihoods (Rigoux et al., 2014; Stephan et al., 2009). This procedure estimates each model’s expected population frequency (model prevalence) and its protected exceedance probability—the probability that it is more frequent than the competing model after accounting for the possibility that model frequencies do not differ.

We validated parameter estimation and model comparison using recovery analyses. For the primary analyses, each model generated one simulated response dataset on each of the 30 participant task sequences in each environment (30 sequences per environment × 2 models × 2 environments = 120 simulated datasets). To concentrate sampling around empirically fitted values while retaining broad coverage, we used a hybrid empirical/broad parameter-sampling scheme: 70% of the generating parameter vectors were sampled intact from participants’ full-data fits, and 30% were obtained by Latin-hypercube sampling across the full fitting bounds listed above, using the same log or logit scales as in fitting. For parameter recovery, we refitted the generating model; all fitted parameters were strongly correlated with their generating values (all Pearson correlations ≥ 0.99; Supplementary Figure 4). For model recovery, we fitted both candidate models to every simulated dataset and repeated the complete cross-validated comparison; the generating model was identified in all 120 cases (Figure 4b). We repeated the recovery analyses using empirical-only and broad-range-only parameter sampling; both showed strong parameter recovery (all Pearson correlations ≥ 0.96) and 100% model recovery, with the generating model correctly identified for every simulated dataset.

Finally, for the fitted-model simulations shown alongside human data, each model was refitted to all 15 blocks from each participant. We simulated that model on the participant’s task sequences using its full-data response-parameter estimates and recomputed the behavioral measures through the same analysis code used for participant responses. These simulations, which served as posterior predictive checks, are shown in the model panels of Figures 5–7, Supplementary Figures 5, 6, and 9, and Supplementary Figure 8.

#### Control and extension models

The **Rescorla–Wagner** control model updates its target paddle-location estimate according to *v_t_*_+1_ = *v_t_* + α δ*_t_*, using the signed prediction error δ*_t_* defined above. Here, α is a constant, latent learning-rate parameter that determines the proportion of the prediction error incorporated into the estimate on each trial. We examined two variants. **Standard Rescorla–Wagner** sets the actual paddle location equal to this target location. **Rescorla–Wagner with response model** instead passes the target location through the same stochastic response model used for the Bayesian models, restricted here to its paddle-location component; this completes the Rescorla–Wagner learner as a model of observed responses by incorporating probabilistic movement initiation and angular response noise.

The formal comparison based on paddle-location responses contrasted two candidates: the environment-matched Bayesian model and Rescorla–Wagner with response model (Supplementary Figure 8, panels e and f). Each candidate had one fitted latent-dynamics parameter alongside the same three fitted location-response parameters: α for Rescorla–Wagner, the change-point probability for the change-point model, and a relative process-to-observation uncertainty scale for the random-walk model. Within each training fold of the five-fold blockwise cross-validation, the latent parameter was selected from a prespecified 37-value grid. The α grid ranged from 0.05 to 0.95 in increments of 0.025. The change-point-probability grid spanned 0.005 to 0.95 with approximately even spacing on the logit scale and included the task-defined probability of 0.0625. Relative uncertainty scales ranged from 0.051 to 4.249 and were chosen to approximately match the steady-state gains corresponding to the α grid. Model recovery applied this complete comparison to 30 synthetic datasets generated by each model in each environment; all 120 datasets were assigned to their generating model. For the fitted-model curves, the models were refitted to all trials and simulated 20 times on each participant’s task sequences. The Standard Rescorla–Wagner curves retained each participant’s fitted α but skipped the stochastic response model, making the target and actual paddle locations identical.

The **hybrid** model runs the change-point and random-walk models in parallel and selects the target response that maximizes expected reward under the weighted mixture of their predictive distributions, *p_mix_*(*x_t_*_+1_| *x*_1:*t*_) = *w p_cp_* (*x_t_*_+1_| *x*_1:*t*_) + (1 − *w*) *p_RW_*(*x_t_*_+1_| *x*_1:*t*_). We fitted the weight *w* separately for each participant using all trials. The fit selected *w* from a 51-value uniformly spaced grid from 0.01 to 0.99 inclusive, alongside the response parameters that entered the fitted likelihood. Values near 1 or 0 approximate the change-point or random-walk model alone, respectively. The primary analysis fitted the model jointly to paddle-location and paddle-width responses. For the component-specific analyses, we refitted the model to paddle-location or paddle-width responses alone, re-estimating the response parameters that entered the corresponding likelihood.

The **stochasticity-decomposition observers** (Supplementary Figure 3) are two simplified variants of the random-walk model that treat volatility as known, with the random-walk SD fixed at the task value of 15°. The first, a Kalman filter, is also given the generative SD (10° or 30°), aligned with trials as in the learning-rate analyses (see *Analysis procedure*). The second infers the generative SD over the same two switching levels as the random-walk model. The theoretical steady-state Kalman gains for fixed process and observation variances are shown as reference lines. For a scalar random walk, these are given by the steady-state solution of the Kalman covariance recursion (Kalman, 1960): 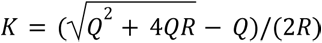, with process variance *Q* = 15^2^ and observation variance *R* = 10^2^ (low variance) or *R* = 30^2^ (high variance).

The **constant-variance model** (Supplementary Figure 10) is a variant of the environment-matched model that assumes a single generative SD, fitted for each participant. To give the variance-inference model fitted variance parameters comparable to those of the constant-variance model, its two SD levels were also fitted, through an overall scale *s* and a high/low ratio *r*, such that σ*_high_* /σ*_low_* = *r* and σ*_high_* σ*_low_* = 300*s*^2^; *s* = 1 and *r* = 3 recover the task values of 10° and 30°. Both models were fitted with the same response model and five-fold blockwise cross-validation as the main comparison, and were compared on the held-out likelihood of paddle-width choices.

#### Implementation

Models were implemented in Python using NumPy and SciPy. The modeling code developed for this study is publicly available in the study’s repository (see *Code and data availability*).

### Data Analyses

#### Apparent learning rate

We quantified learning behavior using the *apparent learning rate*, a model-agnostic measure that has been widely used to characterize human learning (Foucault & Meyniel, 2024; McGuire et al., 2014; Nassar et al., 2010, 2012; Piray & Daw, 2024; Vaghi et al., 2017). For each trial t, the signed paddle-location update was:

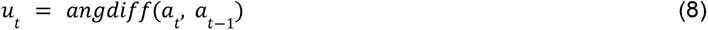

Where *a_t_*_-1_ and *a_t_* are the paddle locations before and after observation *x_t_*, respectively. Using the signed prediction error δ*_t_* defined in *Computational modeling* above, the trial-wise apparent learning rate was:

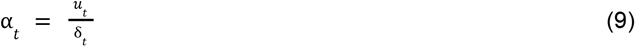

Thus, the update and prediction error were both calculated as signed shortest circular differences. The same calculation was applied to participant responses and simulated model responses. As in previous studies (Foucault & Meyniel, 2024; Piray & Daw, 2024; Vaghi et al., 2017), per-trial apparent learning rates outside the range [−0.6, 1.3] were excluded to limit the influence of outliers; these bounds followed prior work (Foucault & Meyniel, 2024).

#### Analysis procedure

All our behavioral analyses followed a two-step procedure: first computing participant-level measures, then aggregating those measures across participants for the group-level analysis. The same procedure was applied to model simulations: models were run separately for each participant using the exact observation sequences shown in the experiment, enabling the same two-step analysis.

At the participant level, trial-wise apparent learning rates were averaged within bins of absolute prediction error, either across all trials or separately by generative variance. We also averaged apparent learning rates within each variance level without conditioning on prediction-error magnitude. Paddle-width choices (coded 0 for small and 1 for large) were averaged within each variance level. Each variance split used the generative variance most recently in effect when the relevant prediction was made: for apparent learning rates, the variance on the preceding trial, which set the context in which the current prediction error was interpreted; for paddle-width choices, the variance of the observation immediately preceding the choice. A trial was counted as a paddle movement if the paddle location changed. For simulated model responses, changes smaller than 1.96° were counted as no movement, excluding most of the positional noise (circular SD ≈ 1°) that the response model adds when the paddle is not moved. This definition was used both for the proportion of trials with paddle movement and for movement-conditional apparent learning rates.

Event-related analyses were conducted by epoching and averaging paddle-width choices and apparent learning rates within participant around changes in the mean and variance. For the paddle-width duration analysis (Figure 7), the pre-event baseline was the mean across lags −2 and −1, and the post-event peak was the maximum across the first five post-event lags. Duration was defined only when this peak exceeded baseline, and was equal to the number of consecutive observations from the peak for which the curve remained above the value halfway between baseline and peak; a duration of 1 indicated that the curve fell below this halfway value immediately after the peak. The paired comparison included participants for whom duration was defined for both mean change points and variance increases.

#### Statistical analyses

All statistical tests were two-sided unless specified otherwise. Within-environment comparisons used paired or one-sample t-tests; comparisons between the two environments, which comprised different participants, used Welch’s t-tests. Non-parametric analyses used Mann–Whitney, Wilcoxon signed-rank, or binomial tests as reported. For t-tests, effect sizes were Cohen’s d for independent and one-sample comparisons and Cohen’s *d_z_* for paired comparisons; rank-biserial correlations were reported for Mann–Whitney tests.

Three types of comparisons between participant data and fitted-model simulations reported in the Results followed a common participant-level procedure: learning-rate curves across prediction-error bins (Figure 5, Supplementary Figure 6, and Supplementary Figure 8), learning-rate profiles following mean changes (Supplementary Figure 9), and variance-dependent paddle-width adjustments (Figure 6). For each fitted model, we first averaged the relevant measure across 20 independent response simulations for each participant. For the prediction-error and event-related learning-rate curves, we calculated the mean absolute error and mean squared error between the participant and model curves and compared these human–model distances between models using paired t-tests. The prediction-error analysis used raw apparent-learning-rate values across the plotted bins and was repeated for trials in which participants moved their paddle. Event-related curves were baseline-corrected by subtracting the mean apparent learning rate at the two pre-event lags, and distances were calculated across the first eight observations following mean changes. For the paddle-width analysis, we calculated each participant’s high-minus-low change in large-paddle use and its absolute difference from the corresponding change in each model. The resulting distances for the variance-inference model and each fixed-variance control were compared using paired t-tests, with Holm adjustment for the two comparisons within each environment; squared differences and Wilcoxon signed-rank tests were used as sensitivity analyses.

#### Implementation

All analyses were implemented in Python using standard scientific libraries (NumPy, SciPy, pandas, Matplotlib). See the *Code and data availability* section for code access details.

## Code and data availability

All data, code, and results supporting the findings of this study are publicly available on GitHub at https://github.com/cedricfoucault/infer-pred, and archived on Zenodo at https://doi.org/10.5281/zenodo.23263377. The repository allows all results reported in the paper to be fully reproduced. It contains the behavioral data of both experiments, the task code, and the modeling and analysis code, together with the fitted models and the outputs underlying every figure and statistic.

## Acknowledgements

We thank Amy X. Li for her assistance with the ethics approval process. Data collection for this study was funded by a Wellcome/Royal Society Sir Henry Dale Fellowship (208789/Z/17/Z) awarded to L.H., who was also supported by a strategic Longer and Larger award from BBSRC (BB/W003392/1). C.F. was supported by a Junior Research Fellowship from Christ Church, University of Oxford.

## Supplementary Information

**Supplementary Figure 1.**
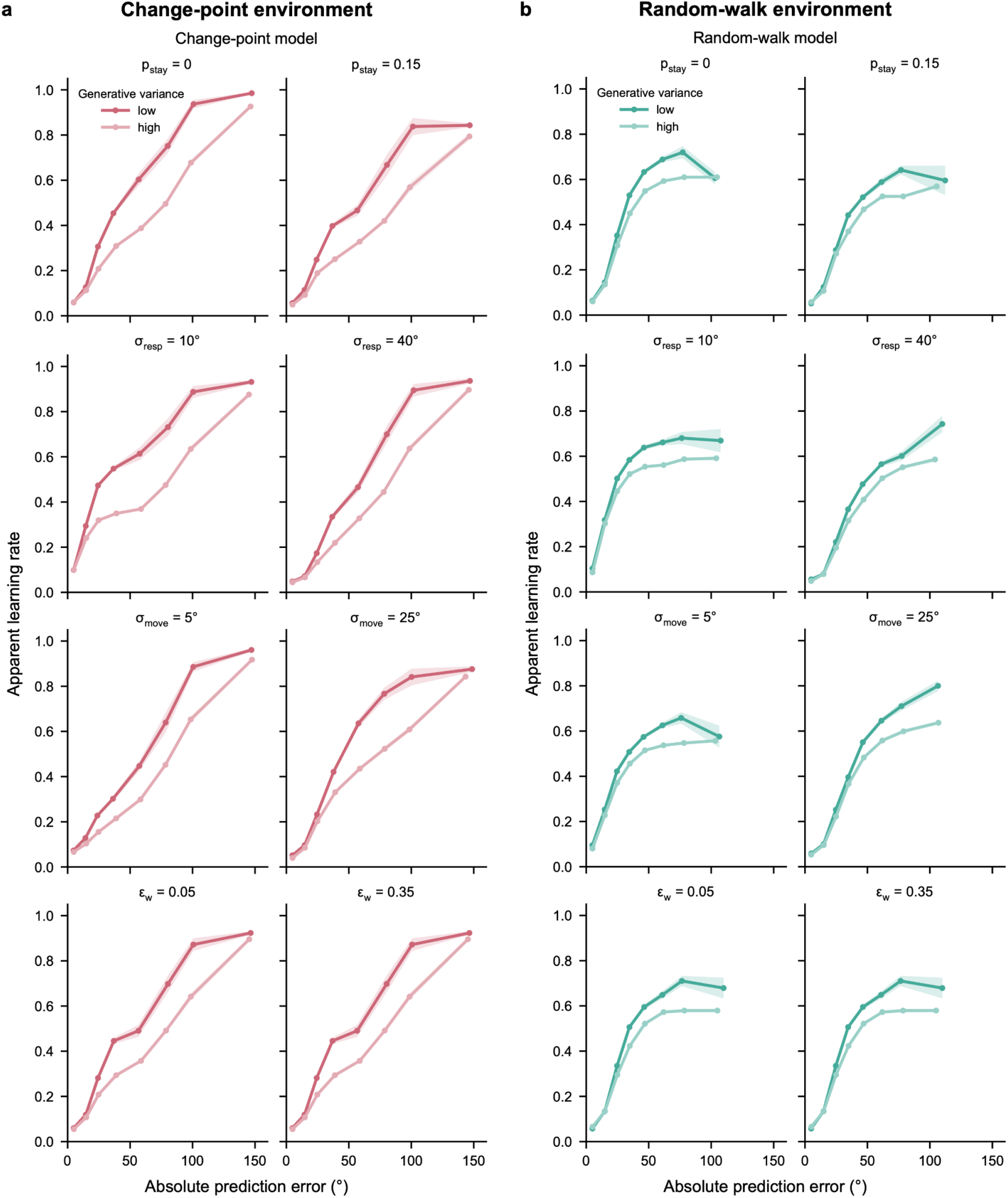
The predicted difference between change-point and random-walk dynamics is robust to the response-model parameters. Apparent learning rate as a function of absolute prediction error, split by generative variance, for the change-point model in the change-point environment (a) and the random-walk model in the random-walk environment (b). Each row shows the effect of one response parameter, set to a low value (left) and a high value (right), with the other parameters held at the canonical values used in Figure 3. In every setting, the apparent learning rate rises more steeply for large prediction errors in the change-point model than in the random-walk model. *ε_w_* affects only paddle-width choices, so its rows match the canonical curves in Figure 3. Shaded areas show ± s.e.m. across simulated participant sequences.

**Supplementary Figure 2.**
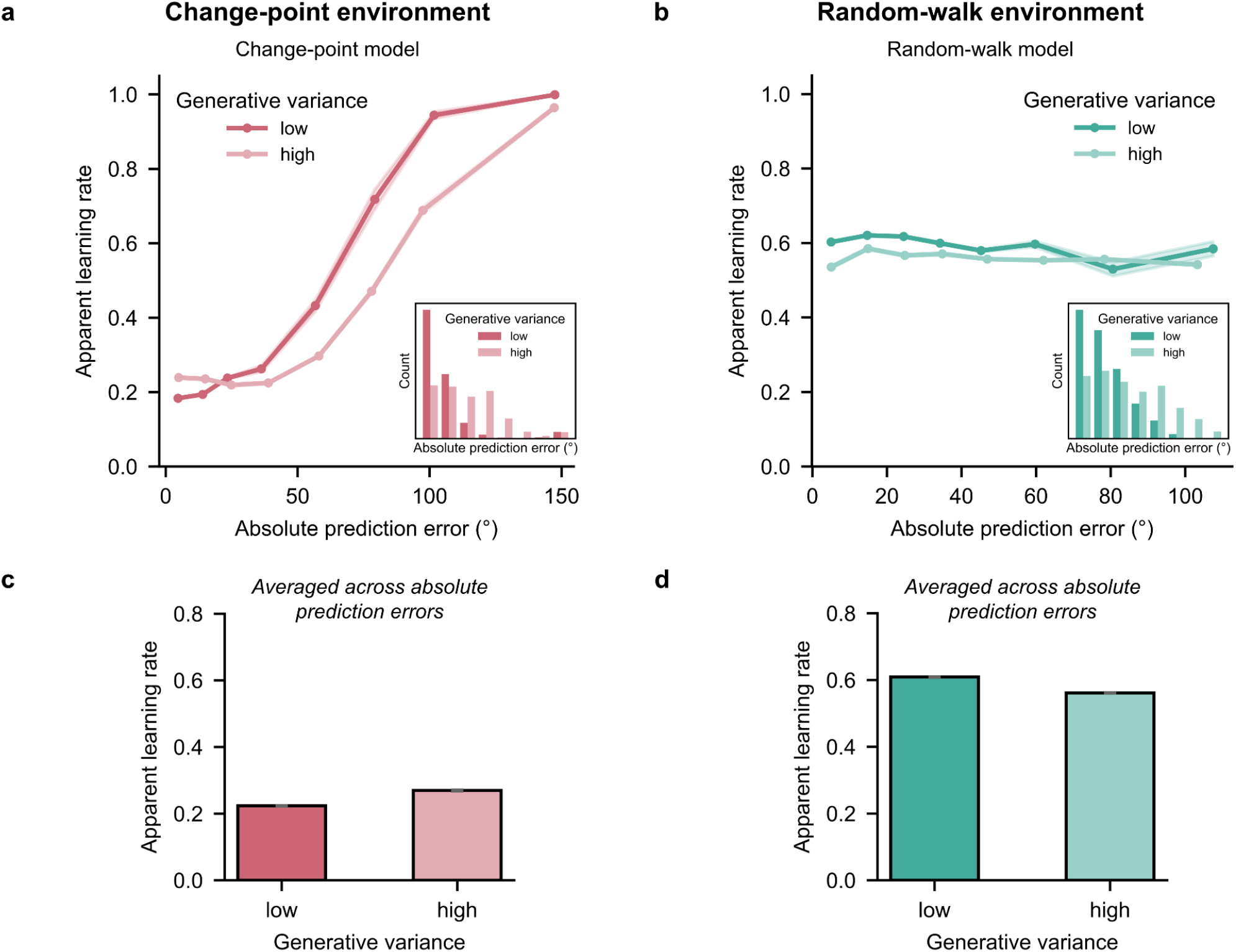
Model predictions without the response model. Same analysis as in Figure 3, except that apparent learning rates are calculated from the models’ target responses—the reward-maximizing responses before the response model is applied (Figure 2a). (a, b) Apparent learning rate as a function of absolute prediction error, split by generative variance, for the change-point model in the change-point environment (a) and the random-walk model in the random-walk environment (b). Insets show the prediction-error distributions. As in Figure 3, apparent learning rate rises more steeply with prediction-error magnitude in the change-point model than in the random-walk model, where it remains nearly constant. Unlike in Figure 3, it is not reduced for small errors, showing that this reduction arises from the response model. (c, d) Mean apparent learning rate under low and high generative variance. The mean is higher under high variance in the change-point model, as in Figure 3c, but lower in the random-walk model, unlike in Figure 3d. Shaded areas and error bars show ± s.e.m. across simulated participant sequences.

**Supplementary Figure 3.**
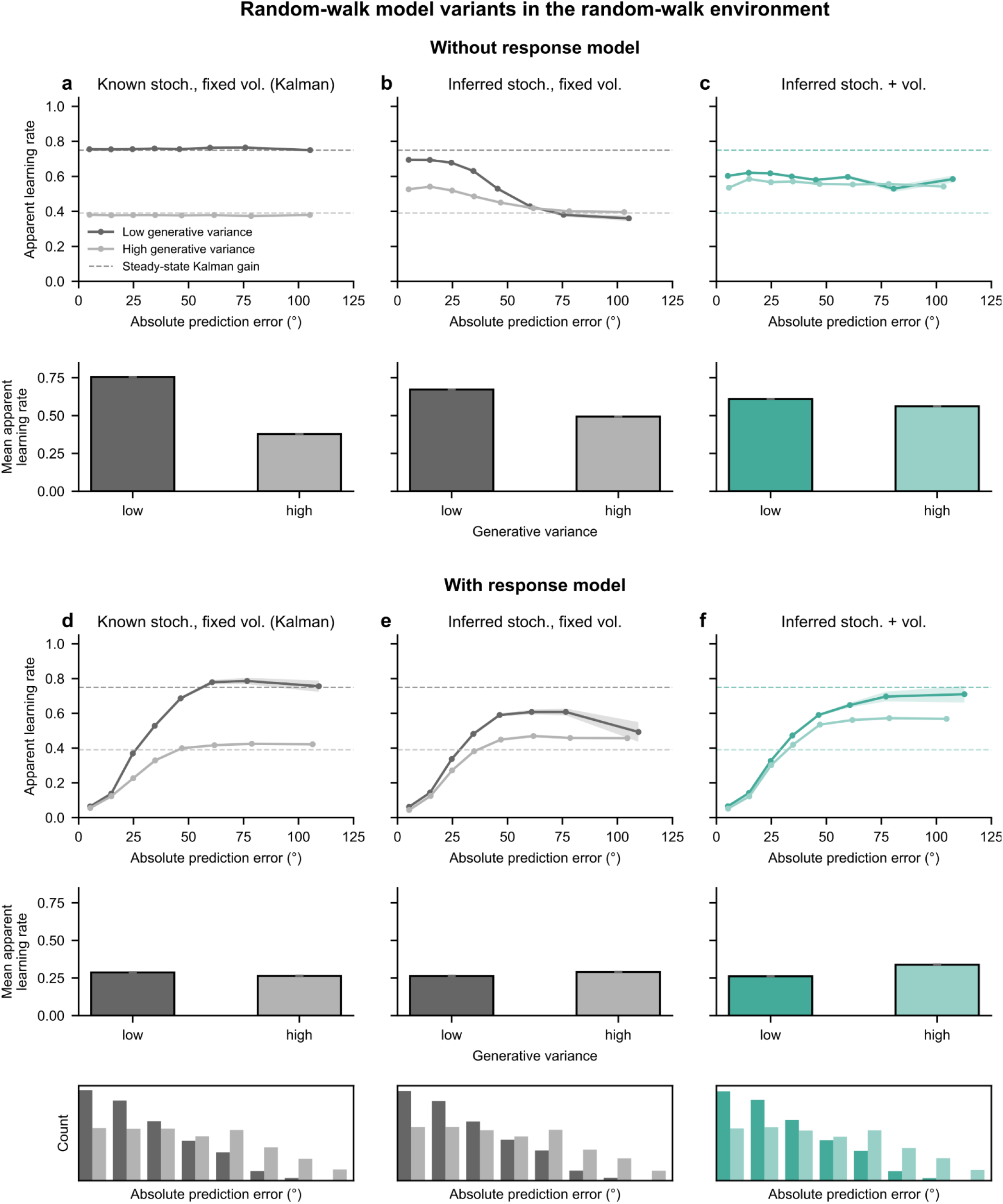
Decomposition of the effects of stochasticity on apparent learning rate. Apparent learning rate of three random-walk observers in the random-walk environment, without (a–c) and with (d–f) the response model. (a, d) The observer is given the stochasticity (generative variance) and assumes a fixed volatility (random-walk SD), which corresponds to a Kalman filter. (b, e) The observer infers stochasticity but assumes the same fixed volatility. (c, f) The observer infers both stochasticity and volatility; this is the random-walk model used in the paper (Supplementary Figure 2b; Figure 3b). Dashed lines mark the theoretical steady-state Kalman gains for low and high variance. Relative to knowing stochasticity (a), inferring it reduces the difference in apparent learning rate between low and high variance (b), because learning rate then follows the inferred rather than the true variance: large errors signal high variance and lower learning rate toward the high-variance gain under both variance levels. Additionally inferring volatility reduces the difference further (c), because noisier observations also increase inferred volatility. With the response model (d–f), mean apparent learning rate can be higher under high variance. Histograms below d–f show the corresponding prediction-error distributions. Response-model parameters, analysis details, and plotting conventions as in Figure 3.

**Supplementary Figure 4.**
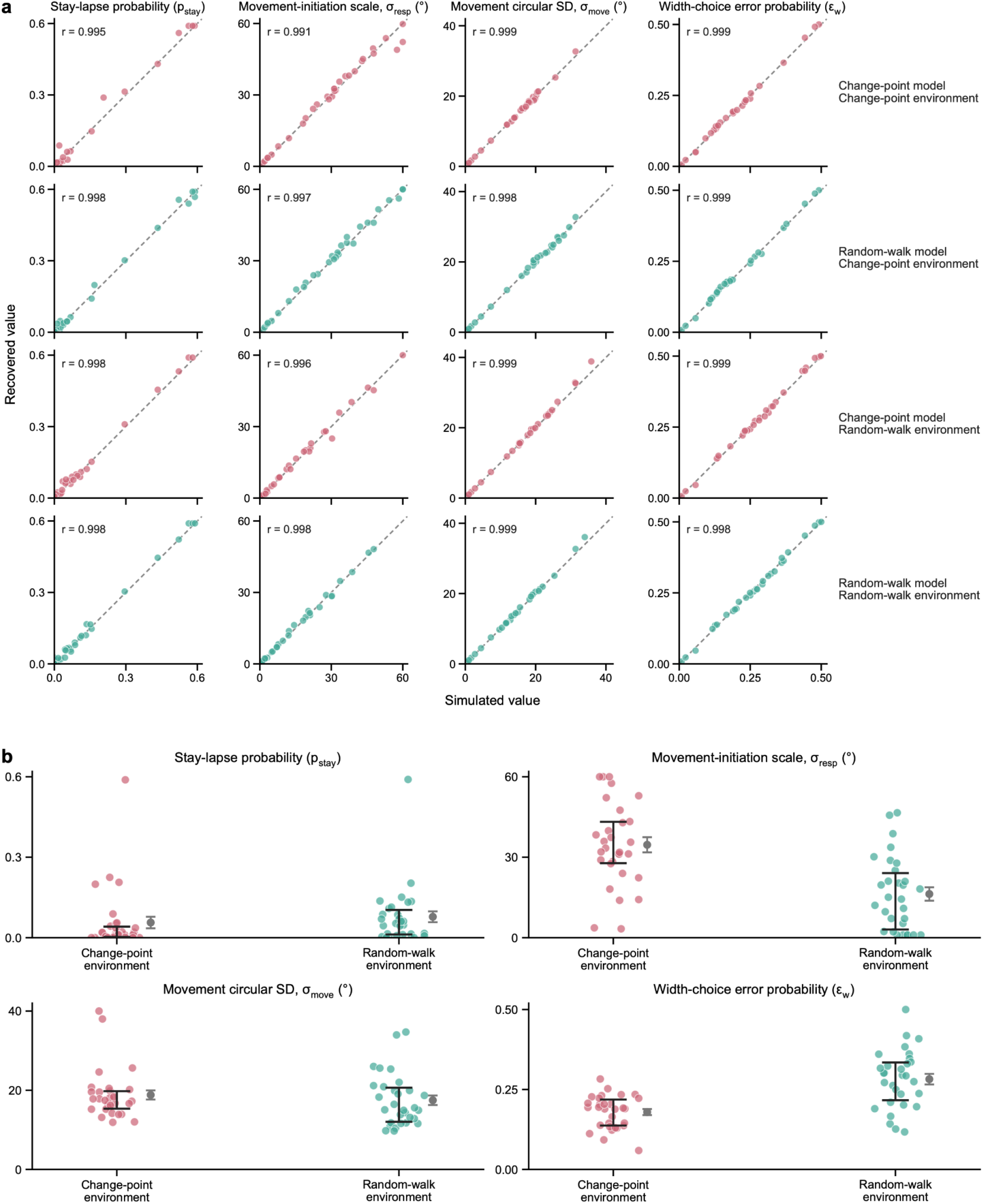
Parameter recovery and fitted response parameters. (a) Recovered versus simulated values of the four response parameters for the change-point and random-walk models in each environment (rows). Each point is one synthetic dataset (30 per model and environment; see Methods); dashed lines show identity. All parameters were accurately recovered (Pearson r ≥ 0.99). (b) Response parameters fitted to each participant’s full data under the environment-matched model. Points show participants; black lines show the median and interquartile range, and gray points and error bars show the mean ± s.e.m.

**Supplementary Figure 5.**
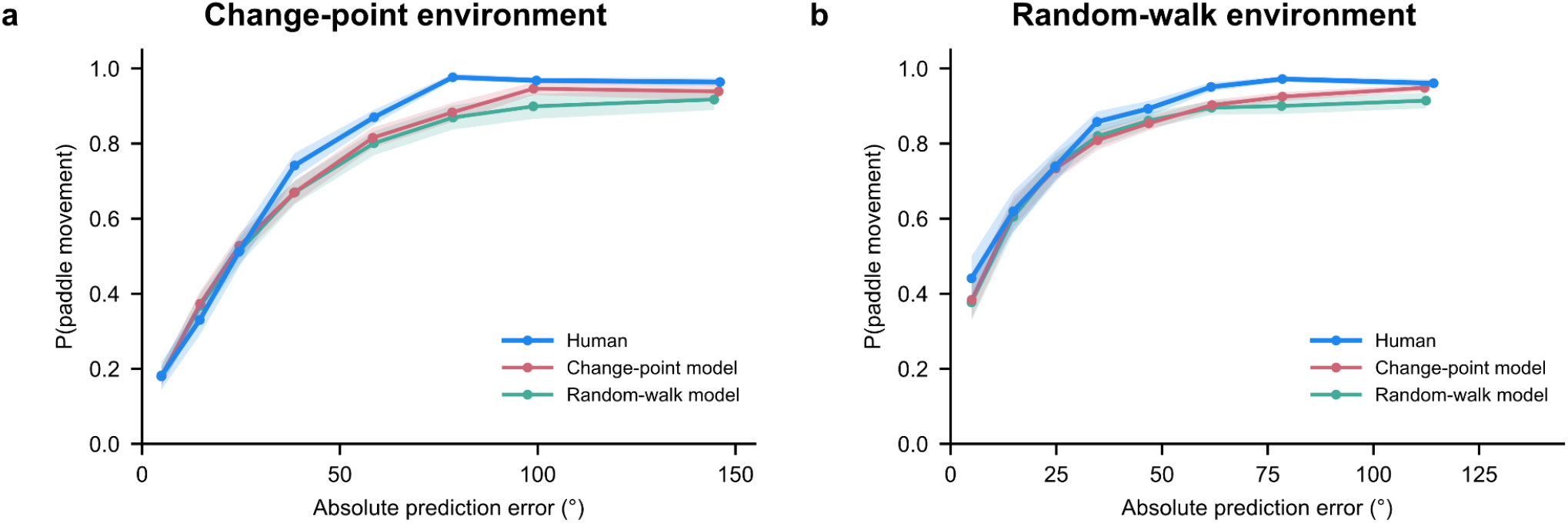
Proportion of trials with paddle movement as a function of absolute prediction error. Human data in the change-point (a) and random-walk (b) environments are overlaid with posterior-predictive simulations of the fitted change-point and random-walk models. Participants were more likely to move the paddle in response to small prediction errors in the random-walk environment than in the change-point environment (statistics in the main text). Both fitted models reproduce this difference through their fitted response parameters, so this measure alone does not distinguish them. Simulation details and plotting conventions as in Figure 5.

**Supplementary Figure 6.**
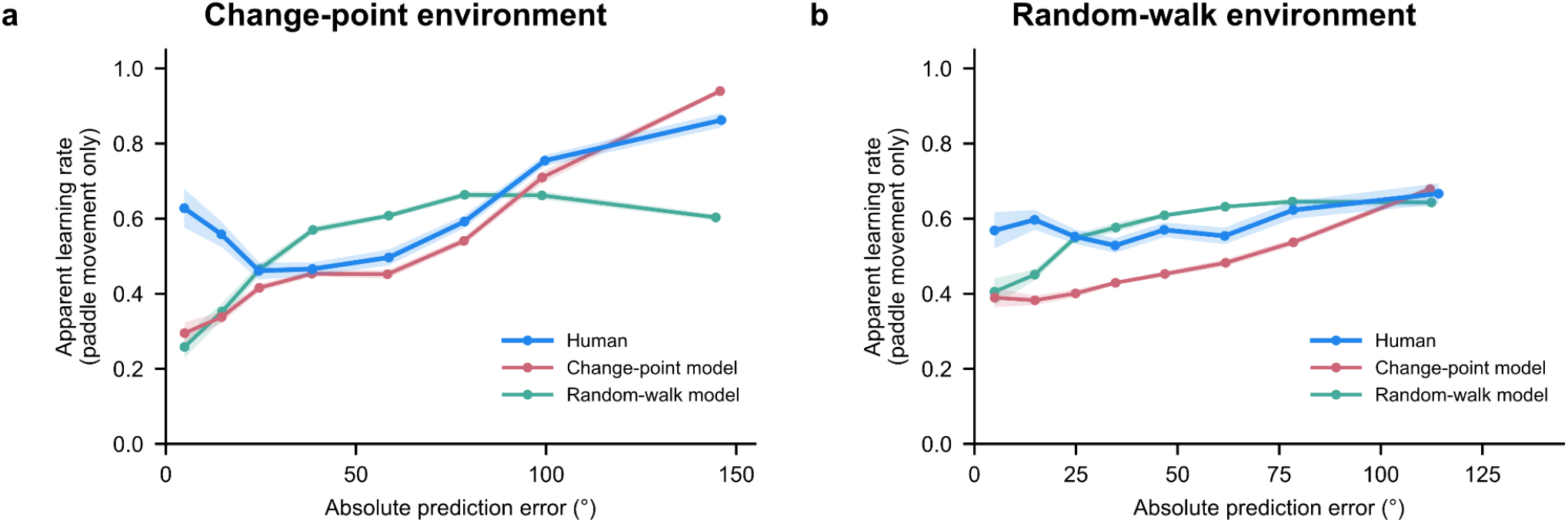
Apparent learning rate on paddle-movement trials as a function of absolute prediction error. Same as Figure 5a, b, except that trials without paddle movement were excluded (see Methods). Human data in the change-point (a) and random-walk (b) environments are overlaid with posterior-predictive simulations of the fitted change-point and random-walk models. When participants moved the paddle, their apparent learning rates still increased more for large prediction errors in the change-point than in the random-walk environment, and in each environment the human curve remained closer to the matched than to the mismatched model (statistics in the main text). This correspondence is therefore not explained solely by how often participants moved the paddle (Supplementary Figure 5).

**Supplementary Figure 7.**
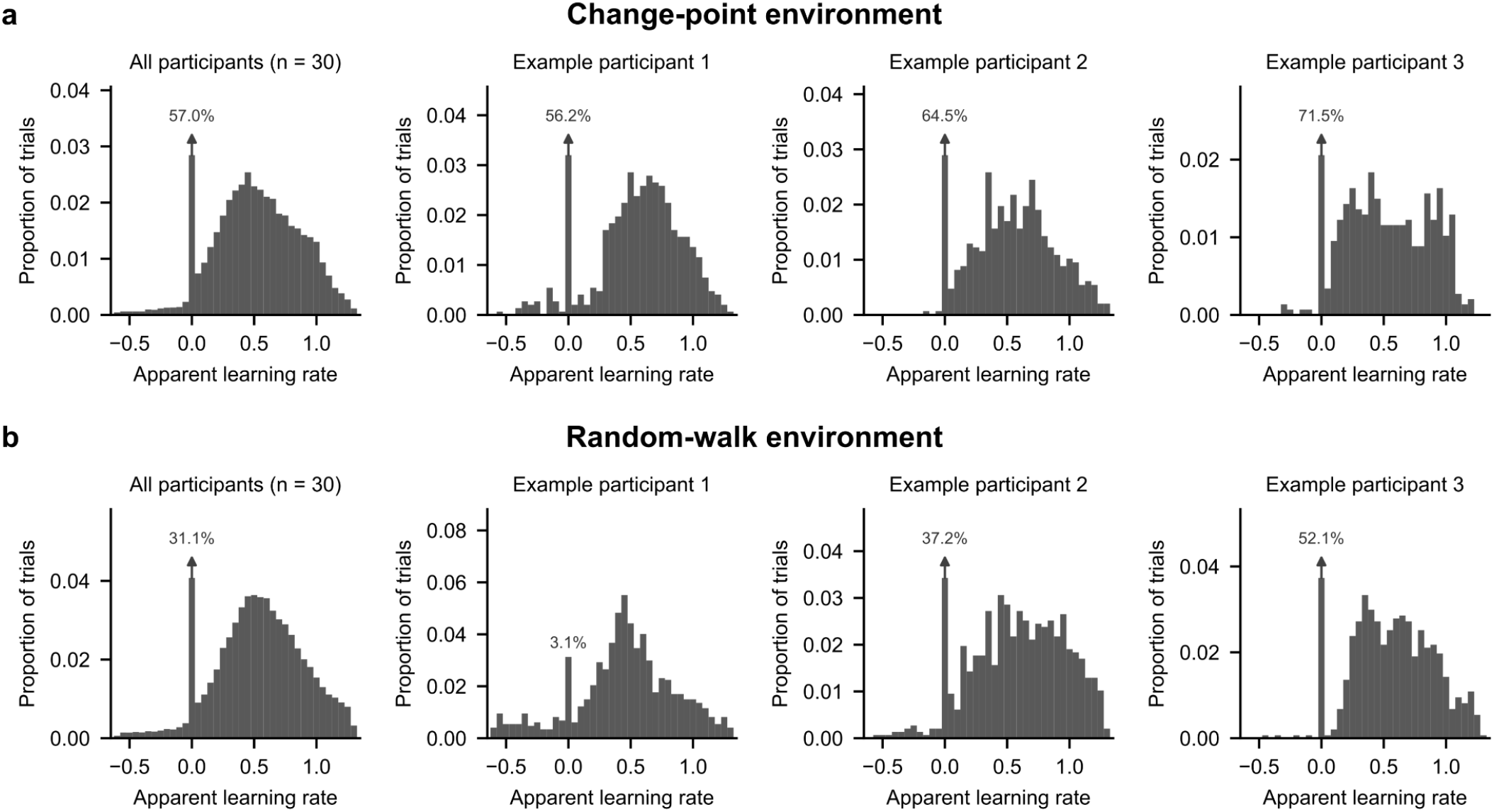
Distributions of trial-wise apparent learning rates. Histograms of human apparent learning rates in the change-point (a) and random-walk (b) environments, pooled across all participants (left) and for three example participants per environment, whose mean apparent learning rates are closest to the 75th, 50th, and 25th percentiles across participants (examples 1–3). The bin at zero, which mainly contains trials without paddle movement, is truncated so that the rest of the distribution remains visible; arrows and labels give its full proportion. Apart from this peak, the distributions show no pronounced accumulation at 1 or at the edges of the plotted range.

**Supplementary Figure 8.**
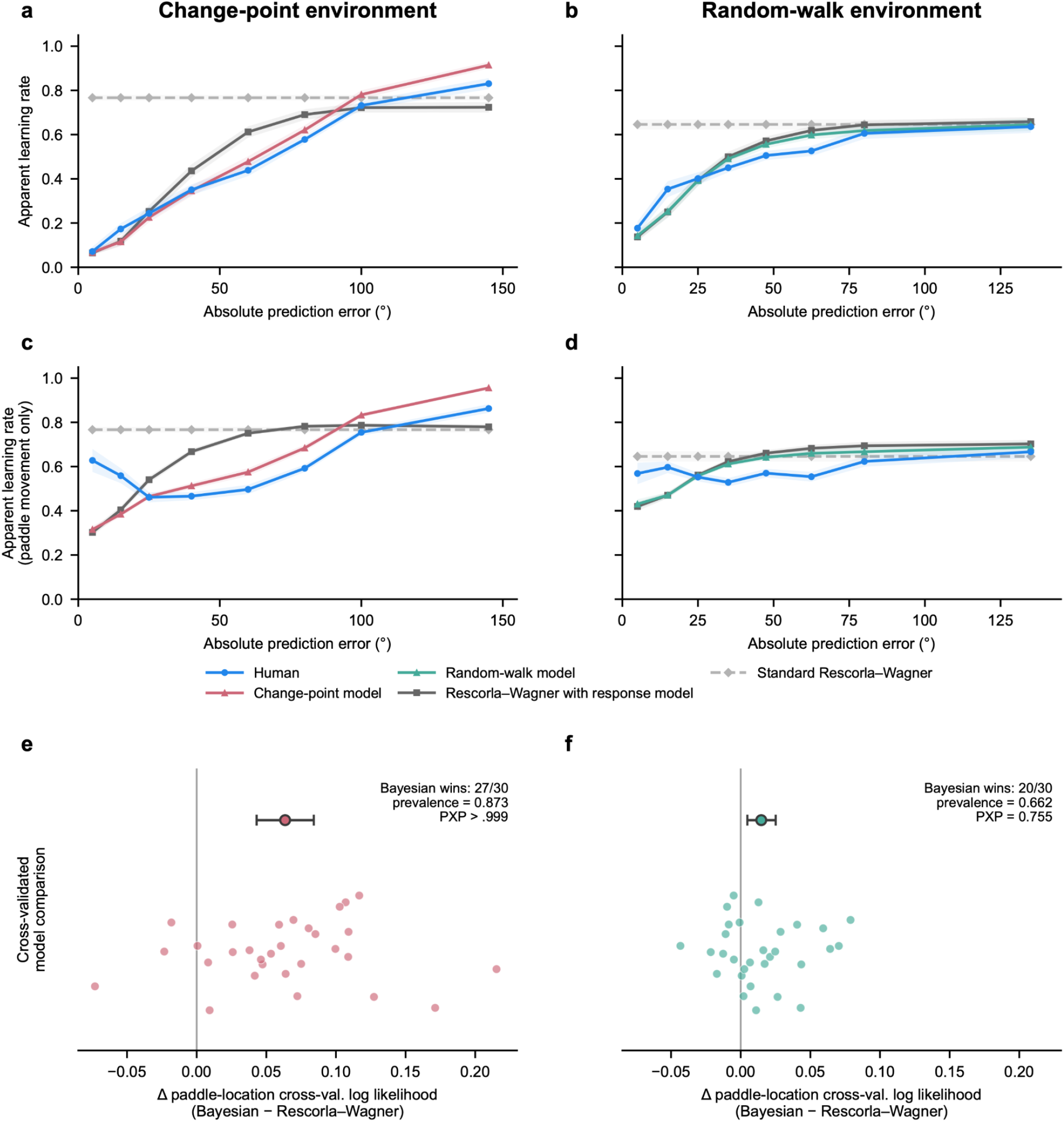
Rescorla–Wagner control models. Human apparent learning rates compared with the environment-matched Bayesian model and a Rescorla–Wagner learner, which updates with a constant latent learning rate, in the change-point (a, c, e) and random-walk (b, d, f) environments. (a, b) Apparent learning rate as a function of absolute prediction error. When passed through the response model, the Rescorla–Wagner learner produces apparent learning rates that rise with prediction-error magnitude, whereas standard Rescorla–Wagner, which uses each participant’s fitted learning rate without the response model, produces a flat curve (dashed). Human curves are nevertheless closer to the Bayesian model than to Rescorla–Wagner with response model (statistics in the main text). (c, d) Same, on paddle-movement trials only. In a–d, model curves show posterior-predictive simulations of models fitted to each participant’s paddle locations (see Methods), and shaded areas show ± s.e.m. across participants. (e, f) Participant-level differences in cross-validated log likelihood of paddle locations between the Bayesian model and Rescorla–Wagner with response model, fitted with the same number of free parameters; positive values favor the Bayesian model. Circles and error bars show the mean and bootstrap 95% confidence interval; annotations give the number of participants favoring the Bayesian model, model prevalence, and protected exceedance probability (PXP).

**Supplementary Figure 9.**
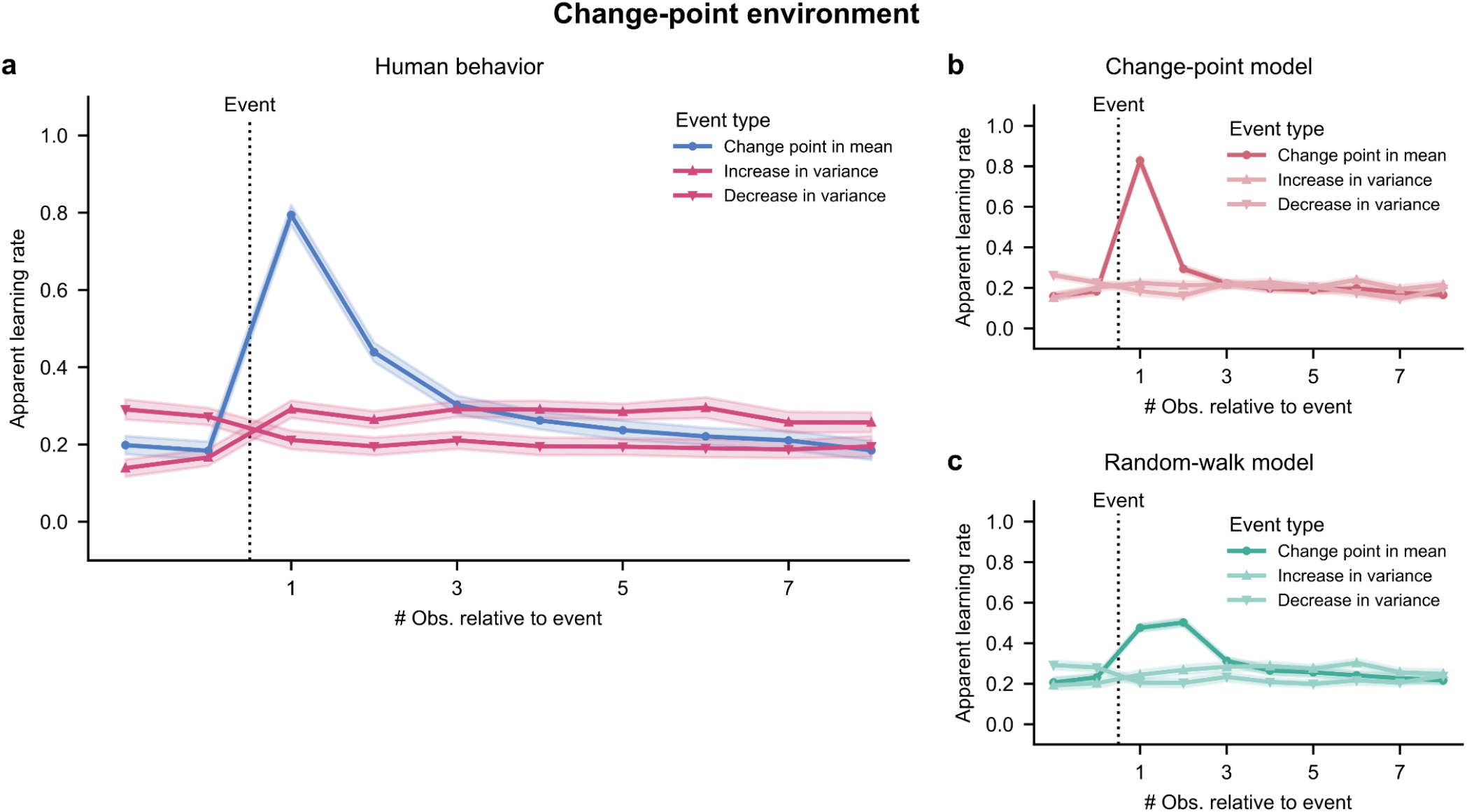
In the change-point environment, human apparent learning rates peak after change points in the mean rather than after changes in variance, as in the change-point model. Apparent learning rate of human participants (a), the fitted change-point model (b), and the fitted random-walk model (c) around change points in the mean, increases in variance, and decreases in variance. Humans and the change-point model show a large transient increase in apparent learning rate at the first observation following a change point in the mean, and at most a small change following changes in variance. The random-walk model shows a smaller increase after mean change points, spread over two observations. The human profile following mean change points is closer to the change-point than to the random-walk model (statistics in the main text). Simulation details and plotting conventions as in Figure 7.

**Supplementary Figure 10.**
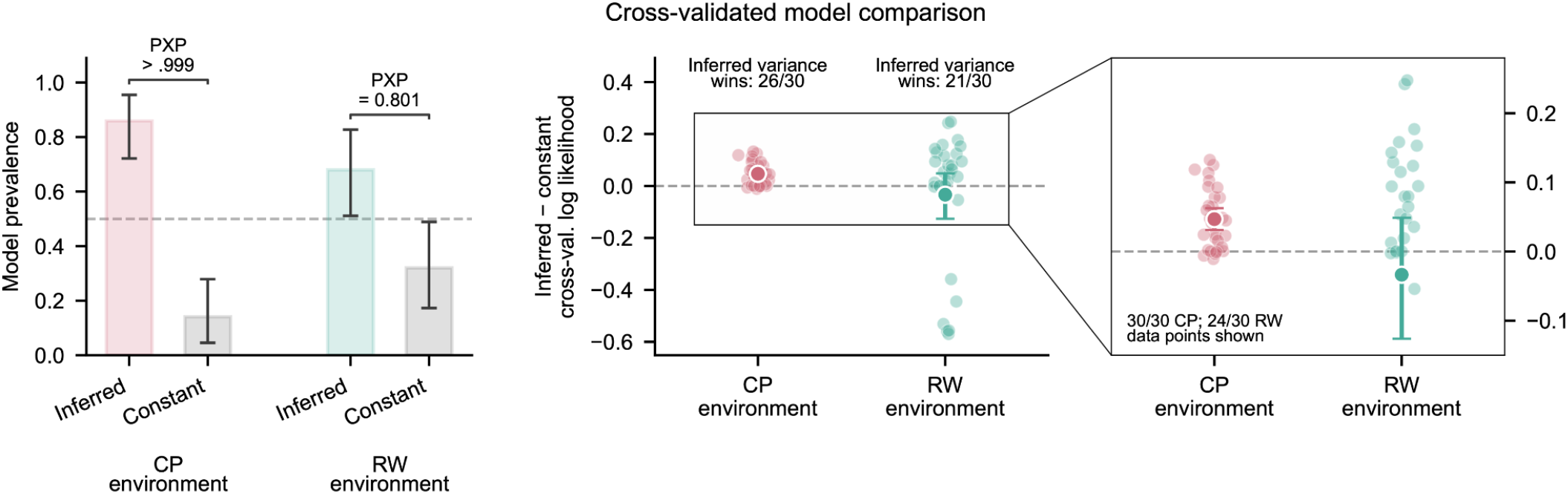
Cross-validated model comparison between variance-inference and constant-variance models. A constant-variance model, which assumes a single fitted variance parameter, was compared with the environment-matched Bayesian model, which infers switches between two variance levels, with comparable fitted variance parameters (overall variance scale and high/low ratio). Both models were fitted to each participant’s responses, and the comparison uses the cross-validated log likelihood of paddle-width choices only. Left: model prevalence from random-effects Bayesian model selection in the change-point (CP) and random-walk (RW) environments. Middle: participant-level differences in cross-validated log likelihood per response between the two models; positive values favor variance inference. Right: enlargement of the framed central range; six RW participants with large negative values lie outside it. Variance inference was favored in most participants in both environments; the evidence was strong in the CP environment and more heterogeneous in the RW environment (statistics in the main text). Plotting conventions as in Figure 4a; means and confidence intervals include all participants.

**Supplementary Figure 11.**
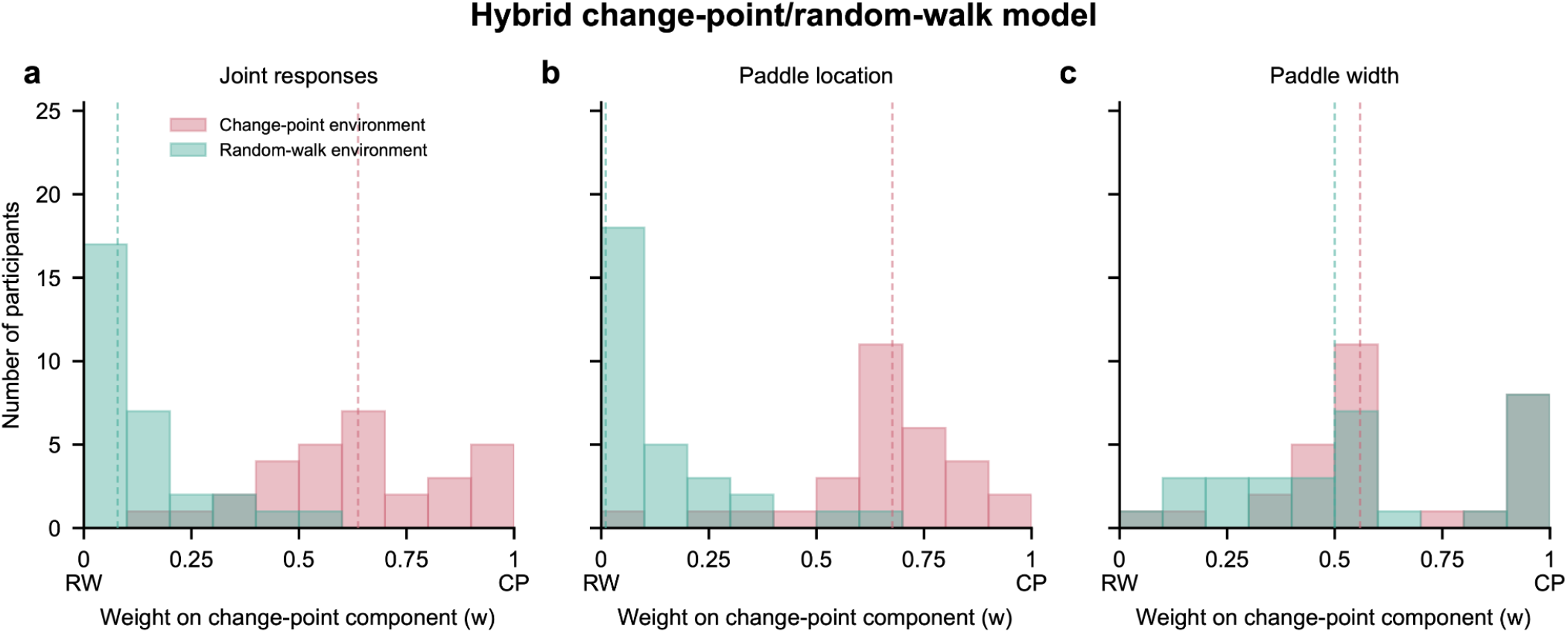
Fitted weights of the hybrid change-point/random-walk model show an environment-dependent shift. Distributions across participants of the fitted mixture weight in the change-point and random-walk environments. The hybrid model was fitted to joint paddle-location and paddle-width responses (a), paddle-location responses only (b), and paddle-width responses only (c). A weight of 1 corresponds to the change-point model and a weight of 0 to the random-walk model. Weights differed strongly between environments for the joint and location-only fits, but not reliably for the width-only fits (statistics in the main text). This pattern is consistent with how the environments and models differ: they differ in the dynamics of the mean, expressed mainly in paddle location, but share the same switching variance, expressed mainly in paddle width. Dashed lines mark the medians. Weights were fitted separately for each participant using all trials (see Methods).

